# Behavioral state and stimulus strength regulate the role of somatostatin interneurons in stabilizing network activity

**DOI:** 10.1101/2024.09.09.612138

**Authors:** Celine M. Cammarata, Yingming Pei, Brenda C. Shields, Shaun S.X. Lim, Tammy Hawley, Jennifer Y. Li, David St. Amand, Nicolas Brunel, Michael R. Tadross, Lindsey L. Glickfeld

## Abstract

Inhibition stabilization enables cortical circuits to encode sensory signals across diverse contexts. Somatostatin-expressing (SST) interneurons are well-suited for this role through their strong recurrent connectivity with excitatory pyramidal cells. We developed a cortical circuit model predicting that SST cells become increasingly important for stabilization as sensory input strengthens. We tested this prediction in mouse primary visual cortex by manipulating excitatory input to SST cells, a key parameter for inhibition stabilization, with a novel cell-type specific pharmacological method to selectively block glutamatergic receptors on SST cells. Consistent with our model predictions, we find antagonizing glutamatergic receptors drives a paradoxical facilitation of SST cells with increasing stimulus contrast. In addition, we find even stronger engagement of SST-dependent stabilization when the mice are aroused. Thus, we reveal that the role of SST cells in cortical processing gradually switches as a function of both input strength and behavioral state.

## Introduction

Normalization is a key function of sensory cortices that allows detection of weak stimuli while preventing saturation to strong stimuli^1–3^. One proposed mechanism for normalization is through amplification of weak inputs via recurrent excitation, which is stabilized by recurrent inhibition as inputs strengthen. Such a network that requires inhibition to avoid runaway excitation is known as an “inhibition-stabilized network” (ISN). A hallmark of an ISN is the paradoxical effect following perturbation of inhibitory interneurons, wherein excitation results in their suppression while suppression yields excitation^4,10–12^. A growing body of work across mice, cats, and primates indicates that auditory, somatosensory, motor and visual cortices exhibit these responses to optogenetic and visual perturbations, suggesting that the cortex generally operates as an ISN^7– 15^. However, it remains poorly understood how the diverse cell types that comprise cortical circuits support inhibition stabilization.

Past research has emphasized the role of parvalbumin-expressing (PV) interneurons in stabilizing network activity^5,10,13,14,16^. These cells receive both feedforward and recurrent excitatory input and robustly inhibit the local excitatory pyramidal cells^22–24^. Empirically, optogenetic stimulation of PV cells yields the hallmark paradoxical suppression^10,13,14^. Moreover, computational modelling has suggested that the PV population is either the exclusive^11^ or the predominant^21^ inhibitory cell type responsible for inhibition stabilization.

Some models, however, indicate that PV cells may be insufficient to stabilize network activity when network excitation is high^25,26^. In such scenarios, network stabilization may additionally require inhibition from somatostatin-expressing (SST) interneurons, which are primarily driven by recurrent excitation from local pyramidal cells and in turn inhibit the pyramidal population^23,27,28^. SST cells are particularly well-positioned to support PV cells in the ISN during high excitation states as they are known to respond robustly to large, high contrast stimuli^16,27,29,30^ and have been implicated in shaping pyramidal output in high arousal states^30–32^. Indeed, optogenetic suppression of SST cells enhances inhibition onto neighboring pyramidal cells, consistent with perturbation of an ISN^13,14^.

Thus, we sought to test whether, and under what network conditions, SST cells are engaged in the ISN. To this end, we developed a model of primary visual cortex (V1) including pyramidal, SST, PV and vasointestinal peptide-expressing (VIP) cells. Our model indicates that while PV cells are initially sufficient to stabilize activity, SST cells are required with increasing sensory input. We tested this prediction in mouse V1 using cell-type specific pharmacology to block AMPA-type glutamate receptors (AMPARs) onto SST cells, thereby selectively reducing the input that connects SST cells to the local network. We find that this manipulation suppresses SST responses to weak visual stimuli, but the suppressive effect is attenuated by strong stimuli or locomotion. Instead, under these conditions, a subset of SST cells is paradoxically driven more strongly following reduction in glutamatergic input. Our computational model reveals that the paradoxical effects that accompany increasing contrast and locomotion are due to the emergence of a network state where stability demands inhibition from SST cells. While the effects of contrast are well-fit solely by increasing input to the network, the effects of locomotion also require changes to local network connections. These results elucidate the conditions under which SST cells are necessary to stabilize visual cortex circuits.

## Results

### A theoretical framework for network stabilization by SST cells

To build an intuition for how input strength and arousal might impact the recruitment of SST cells in stabilizing the network, we developed a model that includes sensory inputs to and connectivity between the four major cortical neuron types: excitatory pyramidal cells (E), and three classes of inhibitory interneurons including SST (S), PV (P), and VIP (V) cells (**Figure 1Ai**). We used a mean-field approach, in which the average firing rate over all cells of a given type is represented by a time-varying scalar (e.g., r_E_ is the average firing rate over all E cells), and each cell type is described by a non-linear input-output transfer function, such as r_E_ = Φ_E_(∑*synaptic input*), which converts synaptic inputs to neural output, ensuring that neural activity cannot be negative. We eliminated six connections known to be weak from the literature^18,22,28^ (*I*_*S*_, *W*_*SP*_, *W*_*SS*_, *W*_*PV*_, *W*_*E V*_, *W*_*VV*_).

**Figure 1.**
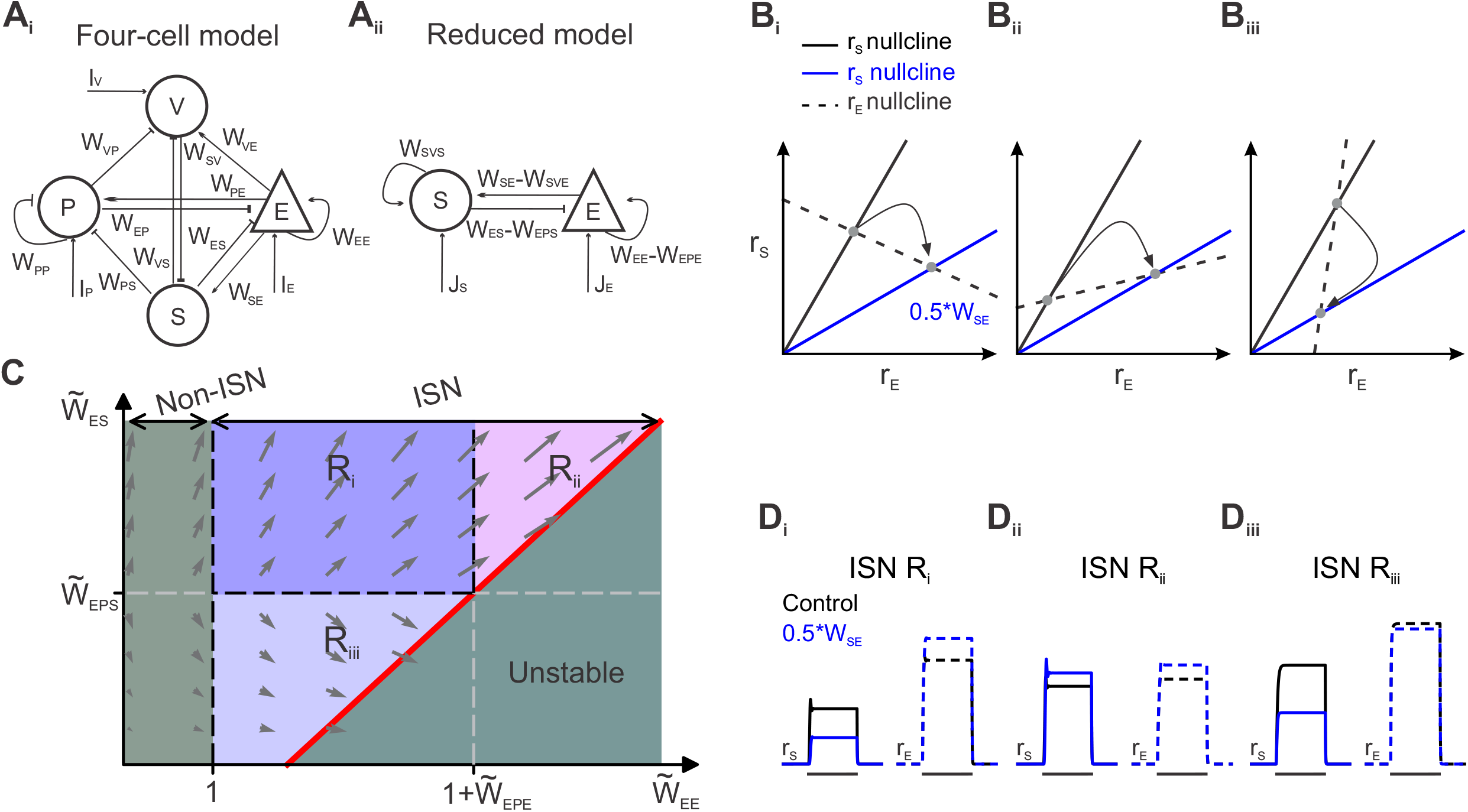
A theoretical framework for network stabilization by SST cells. (A) Schematic of the four-cell (left) and reduced two-cell (right) model. (B_i-iii_) Schematic of r_E_ nullcline (dashed black), r_S_ nullcline in control (solid black) and r_S_ nullcline after a 50% reduction in *W*_*SE*_ (blue) when the slope of the r_E_ nullcline is negative (B_i_), positive (B_ii_) and steeply positive (B_iii_). Arrows illustrate the shift in stability points (gray dots), and therefore the change in r_E_ and r_S_ after decrease in *W*_*SE*_. (C) Network stability in the space defined by 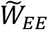 (effective recurrent excitation among E cells) and 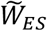 (effective inhibition of S to E). Gray arrows illustrate how effective weights in 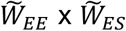 space change when stimulus intensity is increased. (D_i-iii_) Simulated activity of pyramidal (dashed lines) and SST cells (solid lines) in response to a visual stimulus (thick black line) in each region of the space defined in (C) and corresponding to the nullclines illustrated in (B_i-iii_). See also **Figure S1**.

To specifically interrogate the relationship between excitatory pyramidal and SST cells, we reduced the four-cell model to a two-cell model containing only E and S cells (**Figure 1A_ii_**; **STAR Methods**). This two-cell model has four effective synaptic weights, which incorporate the contributions of P and V cells. For instance, the connection from E to E has an effective synaptic weight of *W*_*EE*_ − *W*_*EPE*_, where *W*_*EE*_ is the direct excitatory feedback loop from E to E, while *W*_*EPE*_ reflects an inhibitory feedback loop from E to P back to E (**Figure S1**).

For a given set of effective synaptic weights, the activity of E depends on the activity of S cells (the r_E_ nullcline; **Figure 1B**, dashed line) and vice versa (the r_S_ nullcline; **Figure 1B**, solid line). The intersection of these two lines yields the steady-state activity of E and S cells for the network. Manipulation of the strength of excitation onto S cells (*W*_*SE*_) reduces the slope of the r_S_ nullcline (**Figure 1B**, blue line) and shifts E and S to a new steady state firing rate. When the slope of the r_E_ nullcline is negative, decreasing excitation to S cells results in the expected decrease in S firing rates (**Figure 1B_i_**). However, when the r_E_ nullcline slope is positive, the same manipulation can result in a paradoxical increase in S firing rates, the signature for their requirement for the ISN (**Figure 1B_ii_**). Additionally, when the r_E_ nullcline slope is steeply positive, we find a different paradoxical effect where both E and S rates decrease (**Figure 1B_iii_**). Thus, the necessity of S cells for stabilization depends on the slope of the r_E_ nullcline.

The reduced model further reveals that the slope of the r_E_ nullcline depends on two key parameters: The net recurrent excitation among E cells 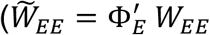, where 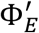 is the derivative of the E current-to-rate transfer function at the current rate) and the net inhibition of S to 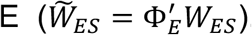.This two-dimensional parameter space has five qualitatively discrete regions: a non-ISN region, three distinct ISN regions (R_i-iii_), and an unstable region (**Figure 1C**), that are defined by four lines. The first line, 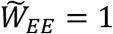, determines whether the network is an ISN (when 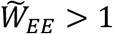) or not, i.e., excitation is weak enough to not require stabilization 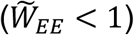.The second line is 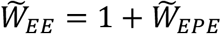, which determines whether the network can be stabilized by P cells alone (when 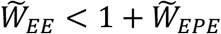). The third line is 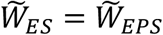, where 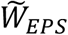 is the strength of S disinhibition of E cells via P (**Figure S1**), which determines whether the net inhibition by S cells outweighs their disinhibition (when 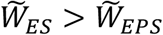). Finally, the fourth line defines the region in which the network is stable (see **STAR Methods**).

The first ISN region (R_i_) is defined by three boundaries: 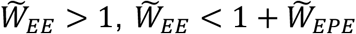, and 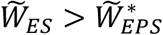. In this region, the network is an ISN, but P cells are sufficient to stabilize the network. In addition, the direct inhibition of E cells by S is stronger than the disinhibition through P cells. In this region, simulating E and S firing rates following a decrease in excitation to S cells (*W*_*SE*_) leads to the intuitively expected result, where S cells have reduced firing rates and E cells are disinhibited (**Figure 1D_i_**).

Starting from region R_i_, increasing 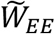 moves the network to the second ISN region (R_ii_) when 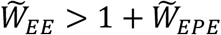. In this region, P cells are no longer able to stabilize the network alone, and thus S cells are also needed for stability. This is revealed by the paradoxical effects of decreasing excitation onto S cells, where like E cells, they increase their firing rates (**Figure 1D_ii_**). Notably, this region in which S cells are required for the ISN is bounded on the high end of 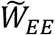 by an unstable region. This boundary is determined by the second axis defined by 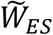. The stronger 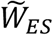, the more 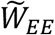 that can be stabilized by S cells. A third ISN region (R_iii_) is defined when 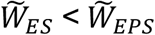.In this region, P cells can stabilize the network alone, but disinhibition of E cells outweighs their direct inhibition from S cells, such that removal of excitation from S cells results in the reduction of both S and E firing rates (**Figure 1D_iii_**). Notably, these three ISN regions define when S cells are necessary, but not when they are sufficient, to stabilize the network (**Figure S1**). Our model makes also predictions about how the network can transition between regions.

First, given that the ISN regimes are defined by synaptic weights, short- and long-term mechanisms that alter synaptic weights^25,34^, such as behavioral state, would be predicted to shift the network state. Second, even if the synaptic weights remain fixed, we predict that network state will be sensitive to input strength. This is because 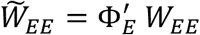 becomes steeper as network activity increases because Φ_E_ is nonlinear. Indeed, simulations of increasing visual stimulus strength move the network from R_i_ towards R_ii_ (**Figure 1C**, arrows). Thus, we predict that SST cells are more likely to be engaged in the ISN with increasing stimulus contrast.

### Cell-type specific antagonism of AMPA receptors

To selectively block excitatory input onto SST cells, and thereby decrease *W*_*SE*_, we used the recently developed cell-type specific pharmacological approach, Drug Acutely Restricted by Tethering^35,36^ (DART; **Figure 2A**). We virally expressed the HaloTag protein (HTP) in V1 of SST::Cre mice to specifically antagonize AMPARs on SST cells upon introduction of YM90K.1^DART.2^ (YM90K^DART^). *In vitro* whole cell recordings reveal that YM90K^DART^ significantly reduces the spontaneous excitatory post-synaptic current (sEPSC) frequency onto HTP-expressing SST cells (^+^HTP cells), compared to control slices in which ^+^HTP cells were incubated in ACSF or a blank^DART^ lacking the YM90K moiety, or the intact YM90K^DART^ applied to SST cells expressing an inactive “double dead” ^dd^HTP (two-way ANOVA main effect for YM90K^DART^, p < 0.001, **Figure 2B-C**). Subsequent application of the traditional AMPAR antagonist NBQX robustly decreases sEPSC frequency onto SST cells in control slices (paired t-test, p = 0.001), but produces only a slight further decrease following YM90K^DART^ (paired t-test, p = 0.106, **Figure 2D**). Interestingly, the amplitude of remaining sEPSCs in the presence of DART is not significantly different from that in control conditions (unpaired t-test, p = 0.117; **Figure 2E**), suggesting a full block at the majority of synapses, rather than fractional block at all synapses. YM90K^DART^ also significantly and specifically reduces the amplitude of electrically evoked EPSCs in SST cells relative to that of concurrently recorded pyramidal cells (**Figure S2A-D**). These data support a specific and robust, but not complete, effect of YM90K^DART^ on SST cells.

**Figure 2.**
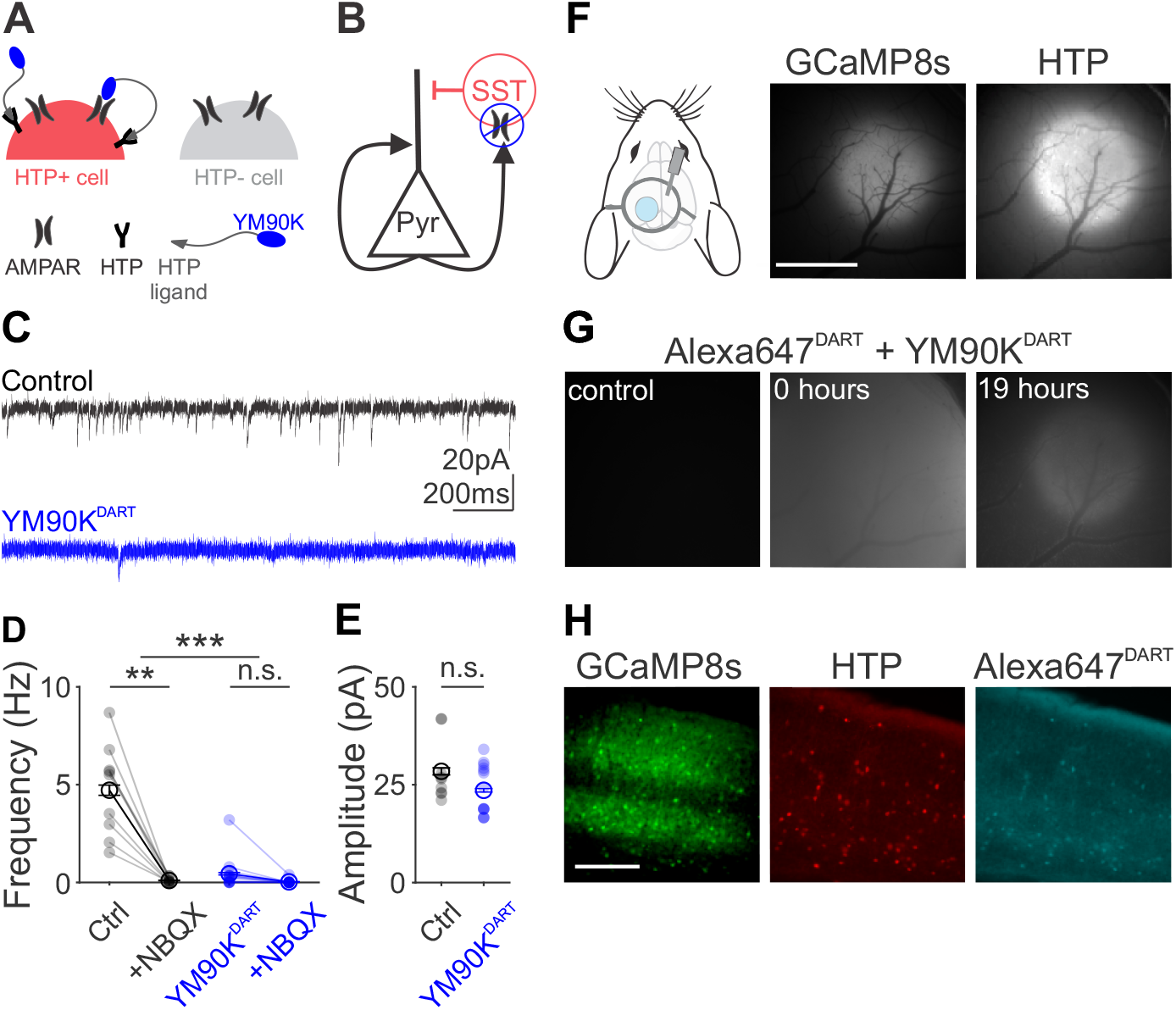
Cell-type specific antagonism of AMPA receptors. (A) Schematic of cell-type specific pharmacology with YM90K^DART^. HTP: Halo-tag protein. (B) Schematic of circuit manipulation. (C) Spontaneous EPSCs (sEPSCs) in an example control SST cell (black) and an example SST cell incubated in YM90K^DART^ (blue). Holding potential is -85 mV to isolate excitatory events. (D) Rate of sEPSCs in normal ACSF or NBQX (10 μM) for control (black) and YM90K^DART^ (blue) cells. Light symbols represent individual cells; dark symbols represent the mean; lines connect individual cells. Error is SEM across cells. (E) Same as (D), for sEPSC amplitude in normal ACSF. (F) Schematic of cranial window and infusion cannula (left), and widefield imaging of the calcium indicator GCaMP8s (middle) and flex-dTomato-HTP (right). Scalebar = 1 mm. (G) Alexa647^DART^ (1:10 with YM90K^DART^) capture before (left), immediately after (middle) and 19 hours after (right) infusion for mouse in (F). (H) Expression of GCaMP8s (left) and HTP (middle), and capture of Alexa647^DART^ (right) in coronal sections for the same mouse as (F-G). Scalebar = 200 µm. n.s.-not significant; ** p < 0.01; *** p< 0.001. See also **Figure S2**.

To probe the effects of blocking excitatory input to SST cells *in vivo*, we pan-neuronally expressed GCaMP8s alongside cell-type specific expression of HTP in V1, and delivered DART ligands via a cannula in the contralateral ventricle (**Figure 2F**). Co-infusion of a mixture of YM90K^DART^ and Alexa647^DART^ enables visualization of the efficacy of ligand delivery and subsequent capture through the cranial window^36^(**Figures 2G** and **S2**). *Post-hoc* histology reveals robust and selective ligand capture on ^+^HTP cells (**Figure 2H**).

### The effect of blocking AMPARs on SST cells depends on stimulus strength

We used two-photon excitation of GCaMP8s to monitor the activity of populations of ^+^HTP SST cells and neighboring putative pyramidal cells in layer 2/3 of V1 while mice passively viewed full-field sinusoidal gratings (2 Hz, 0.1 cycles per degree) moving in one of eight directions (45° increments) at one of three contrasts (25%, 50%, and 100%; **Figure 3A**). Data were collected from the same neurons on consecutive days to measure visual responses in control conditions and 17-24 hours after infusion of YM90K^DART^ (**Figure 3B**).

**Figure 3.**
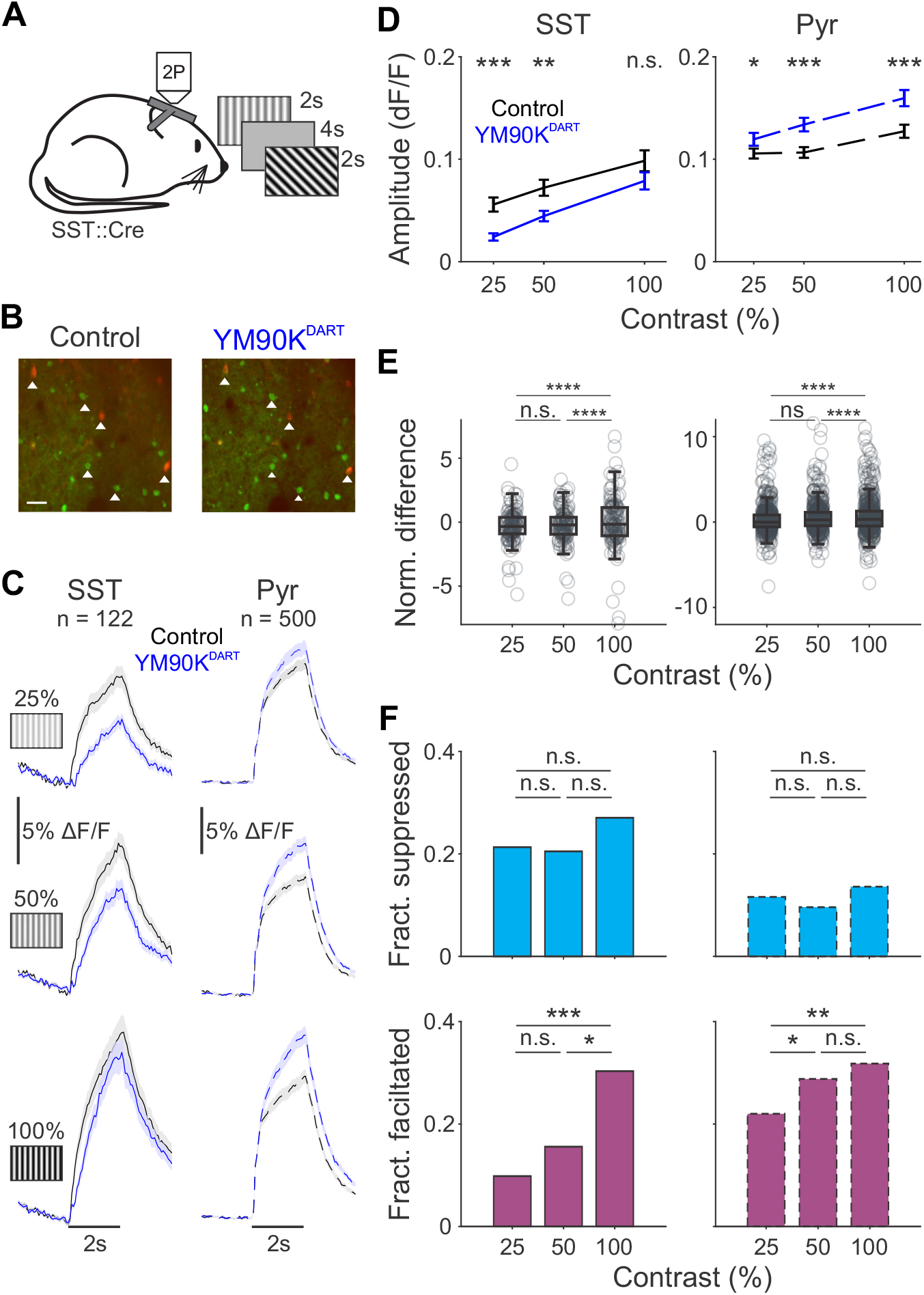
The effect of blocking AMPARs on SST cells depends on stimulus strength. (A) Schematic of experimental setup. (B) Example two-photon imaging field of view of GCaMP (green) and HTP (red) expression in control (left) and after YM90K^DART^ infusion (right) for the same mouse as **Figure 2F-H**. White triangles highlight example cells identifiable across sessions. Scalebar = 200 µm. (C) Grand average time courses for HTP+ SST (left, solid lines) and HTP-putative pyramidal cells (right, dotted lines) before (black) and after (blue) YM90K^DART^ infusion, in response to preferred-direction gratings (horizontal black bar) at three stimulus contrasts, during stationary epochs. Shaded error is SEM across cells. (D) Mean response during stimulus period, for SST cells (left) and pyramidal cells (right) before (black) and after (blue) YM90K^DART^ infusion, at each contrast. Error is SEM across cells. (E) Normalized difference 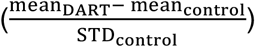 of stimulus response for SST (left) and pyramidal cells (right) as a function of contrast. Gray circles are individual cells; box plots illustrate median, 25% and 75% quartiles. Significance refers to pairwise F tests for variance. (F) Fraction of SST (left) and pyramidal (right) cells that are suppressed (top, cyan) or facilitated (bottom, magneta) by more than 1 std of their control response at each contrast. n.s.-not significant; * p < 0.05; ** p < 0.01; *** p< 0.001; **** p< 0.0001. See also **Figure S3**.

Consistent with the predictions of our model (**Figure 1C**), we find that the magnitude of the effect of blocking AMPARs on SST cells depends on the strength of the visual stimulus. When mice are stationary and the stimulus contrast is low, the population of SST cells has a decreased visual response following YM90K^DART^ (n = 122 cells, 10 mice; paired t-test with Bonferroni correction, p < 0.001, **Figure 3C-D**), consistent with the decrease of excitatory drive. However, with increasing contrast, the effect of DART on the response of SST cells is diminished, such that there is no significant effect of YM90K^DART^ at full contrast (paired t-test with Bonferroni correction, p = 0.203). We do not think that this stimulus dependence is due to elevated glutamate release outcompeting YM90K^DART^ because if this were the case, then we would expect pyramidal cells to exhibit a similar contrast-dependent decrease in effect size. Contrary to this, we find that the effect on the pyramidal cell population increases with increasing contrast (n = 500 cells; two-way ANOVA, interaction of contrast and YM90K^DART^, p = 0.001, **Figure 3C-D**). This argues that the network effects of YM90K^DART^ are actually more robust at high contrast, despite the apparent decreased effect on the average response of SST cells.

To understand why the average effects on SST cells decrease, we investigated the effects on individual cells. We find that the activity of individual SST cells is more strongly modulated by YM90K^DART^ with increasing contrast (Levene’s test for unequal variance, p = 0.001; **Figure 3E**). This is due to the fraction of SST cells that are significantly facilitated by YM90K^DART^ (defined as the mean response increasing more than one standard deviation from control) becoming greater with increasing contrast (chi-square test with Bonferroni correction for 25% vs. 50%, p = 0.535, 25% vs. 100%, p < 0.001; 50% vs. 100%, p = 0.018, **Figure 3F**). This mirrors the increased fraction of pyramidal cells facilitated with greater contrast (25% vs. 50%, p = 0.041; 25% vs. 100%, p = 0.001, 50% vs. 100%, p = 0.960), consistent with inhibition stabilization. In comparison, we find no significant change in the fraction of SST cells that respond with simple suppression (decreased by more than one standard deviation from control: chi-square with Bonferroni correction p > 0.05 for all contrast comparisons), and only a small fraction of pyramidal cells are suppressed at any contrast. Thus, we observe diverse effects on individual SST cells with some being suppressed but more being facilitated as contrast increases.

As a control for ambient-drug effects of YM90K^DART^ and habituation that may occur with repeated imaging^37,38^, we repeated the experiment with YM90K^PEG^ which is chemically identical except for its lack of an HTP ligand. This construct, which cannot bind to HTP, washes out by the time of imaging (n = 6 mice; **Figure S2E-F**). Unlike the effects of YM90K^DART^, treatment with YM90K^PEG^ results in weak suppression of both SST (n = 84 cells; two-way ANOVA, main effect of YM90K^PEG^, p = 0.001; **Figure S3A-D**) and pyramidal (n = 458 cells; p = 0.003) responses, without contrast dependence (two-way ANOVA, contrast x YM90K^PEG^ interaction in SST cells, p = 0.194). Thus, the observed contrast-dependent effects of YM90K^DART^ are due to its action on SST cells.

Together, these findings suggest that reducing excitatory input on SST cells largely results in a straightforward decrease in SST responses at low contrast, but at higher contrast paradoxically increases the visually evoked responses in a subset of SST cells. Stronger visual input results in more robust disinhibition of pyramidal cells, driving the SST cells more strongly via their remaining unblocked glutamate receptors, and ultimately resulting in a net facilitation of their activity. This is consistent with SST cells being recruited to stabilize network activity as stimulus strength increases, as predicted by our theoretical model.

### SST cells correlated with the local network are less suppressed by YM90K^DART^

In an ISN, recurrent input from pyramidal cells recruits interneurons to stabilize the network^4–7^. Given the importance of this recurrent connection for engagement in an ISN, those SST cells that are most robustly recurrently connected should be the most susceptible to paradoxical effects.

Noise correlations can be used as a proxy for shared connectivity^39,40^. Thus, to estimate the strength of recurrent input onto each SST cell, we calculated the noise correlation between individual SST cells and the mean of all simultaneously recorded pyramidal cells during the control imaging session (**Figure 4A**). Correlation measures were pooled across stimulus conditions as we find no significant dependence on contrast (one-way ANOVA, p = 0.055) consistent with past reports^41^. This yields a range of correlation values across the SST population that we defined as weakly (R < 0.5; n = 67 cells; **Figure S4A**) or strongly correlated (R > 0.5; n = 55 cells), with approximately half the SST cells in each experiment falling into each category (fraction strongly correlated, mean across mice ± standard deviation = 46.67% ± 21.37%). We posit that SST cells that are more strongly correlated with pyramidal cells are likely to be more strongly recurrently connected, and therefore less suppressed by YM90K^DART^.

**Figure 4.**
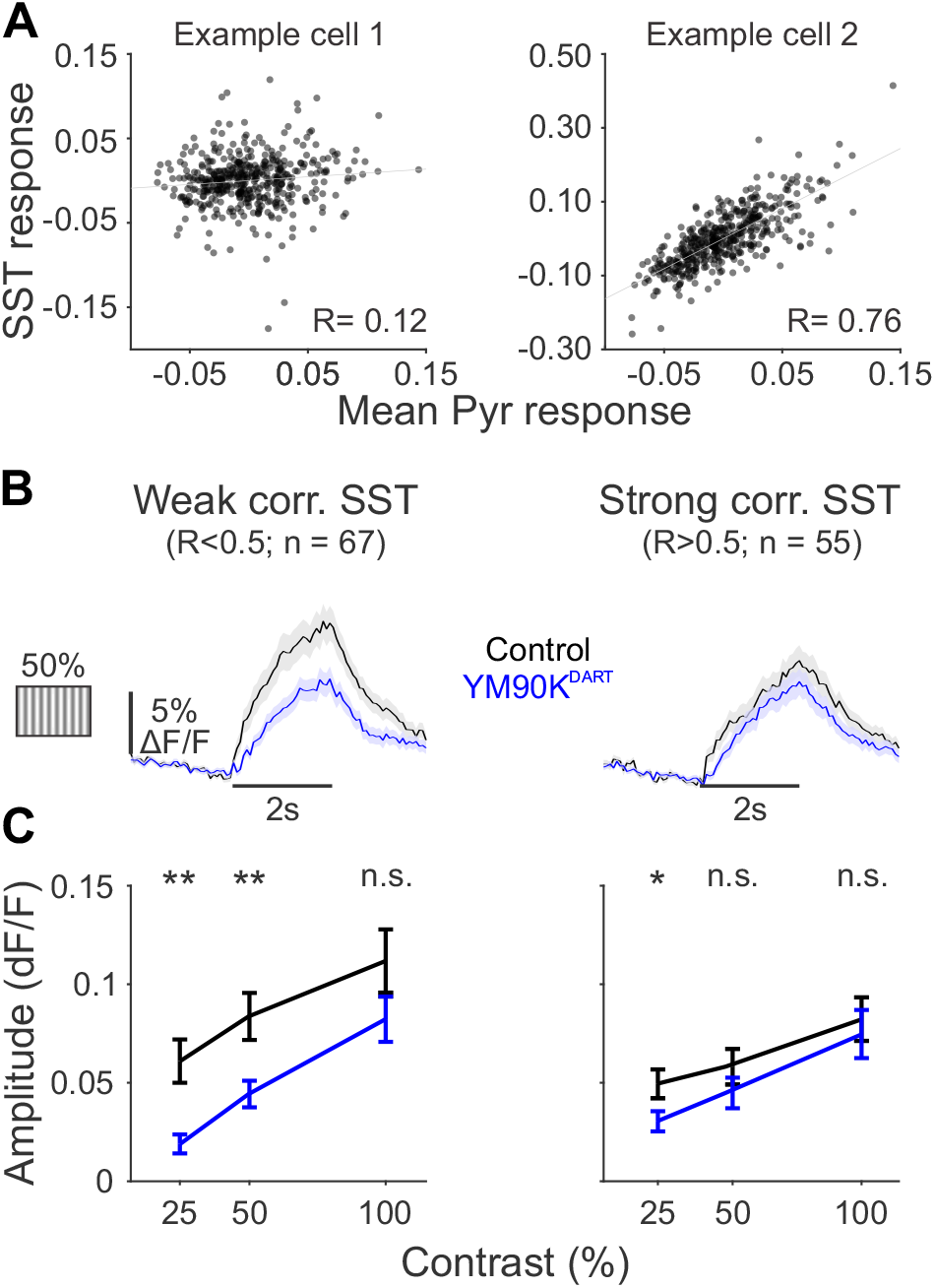
SST cells weakly correlated with the local network are more strongly suppressed by YM90K^DART^. (A) Mean-subtracted trial-by-trial responses for two example SST cells and all concurrently recorded pyramidal cells. Each data point represents a single trial. Fit line is from a linear regression; R is the Pearson’s correlation. (B) Grand average time courses for SST cells before (black) and after (blue) YM90K^DART^ separated into those weakly (left) and strongly (right) correlated to pyramidal activity, during stationary epochs in response to preferred-direction gratings at 50% contrast. Shaded error is SEM across cells. (C) Mean response during stimulus period, for SST cells weakly (left) or strongly (right) correlated to pyramidal activity, at each contrast in control (black) and after YM90K^DART^ (blue). Error is SEM across cells. n.s.-not significant; * p < 0.05; ** p < 0.01. See also **Figure S4**.

Consistent with our prediction, following YM90K^DART^ delivery, the weakly correlated SST cells have a significant decrease in visually evoked responses (two-way ANOVA, main effect of YM90K^DART^, p = 0.003; **Figure 4B-C**), whereas the strongly correlated SST cells are not significantly affected (p = 0.113). This dependence on correlated variability is robust to resampling within, but not across, correlation groups (**Figure S4B-C**), and is specific to YM90K^DART^, as YM90K^PEG^ weakly suppresses both groups (two-way ANOVA, main effect of YM90K^PEG^: weakly correlated cells, n = 48, p = 0.048; strongly correlated cells, n = 36, p = 0.004, **Figure S4D-F**). These results suggest that recurrent excitation determines the effect of YM90K^DART^ on SST cells, consistent with recruitment of SST cells into an ISN. In addition, these result hint that there may be some functional heterogeneity among the population of SST cells that impacts their engagement in the ISN.

### The effect of blocking AMPARs on SST cells depends on behavioral state

Having determined that strong sensory input recruits SST cells to stabilize network activity, we wondered whether other conditions that increase excitation in the V1 cortical network would have a similar effect. Locomotion is well known to increase firing rates in V1^32,34,42^. To compare the same cells across behavioral states, we examined the subset of SST and putative pyramidal cells which could be measured at their preferred direction, in both stationary and running conditions, for all contrasts, and during both imaging sessions. Due to variation in animals’ tendency for running, this led to the exclusion of two mice from both the YM90K^DART^ (n = 8 mice, 91 SST and 379 pyramidal cells; **STAR Methods**) and YM90K^PEG^ (n = 4 mice, 54 SST and 275 pyramidal cells) experiments.

Consistent with previous reports, we find that both SST (three-way ANOVA, main effect for locomotion, p < 0.001; **Figure 5A-B**) and pyramidal cells (p < 0.001; **Figure 5C-D**) are robustly facilitated by running. Moreover, we find that locomotion dramatically changes the impact of YM90K^DART^ on SST cells (three-way ANOVA, YM90K^DART^ x locomotion interaction, p = 0.020; **Figure 5A-B**). The straightforward suppression observed at low contrast when the mice are stationary (paired t-test with Bonferroni correction, p = 0.003) no longer occur when mice are running (p = 0.824). At high contrast, the average response of SST cells trends towards the paradoxical elevation expected from inhibition stabilization, although this did not reach significance (p = 0.253). Moreover, these effects are specific to the block of AMPARs on SST cells, as there is no dependence of the effects of YM90K^PEG^ on behavioral state (three-way ANOVA, YM90K^PEG^ x locomotion interaction, p = 0.488, **Figure S3A-D**).

**Figure 5.**
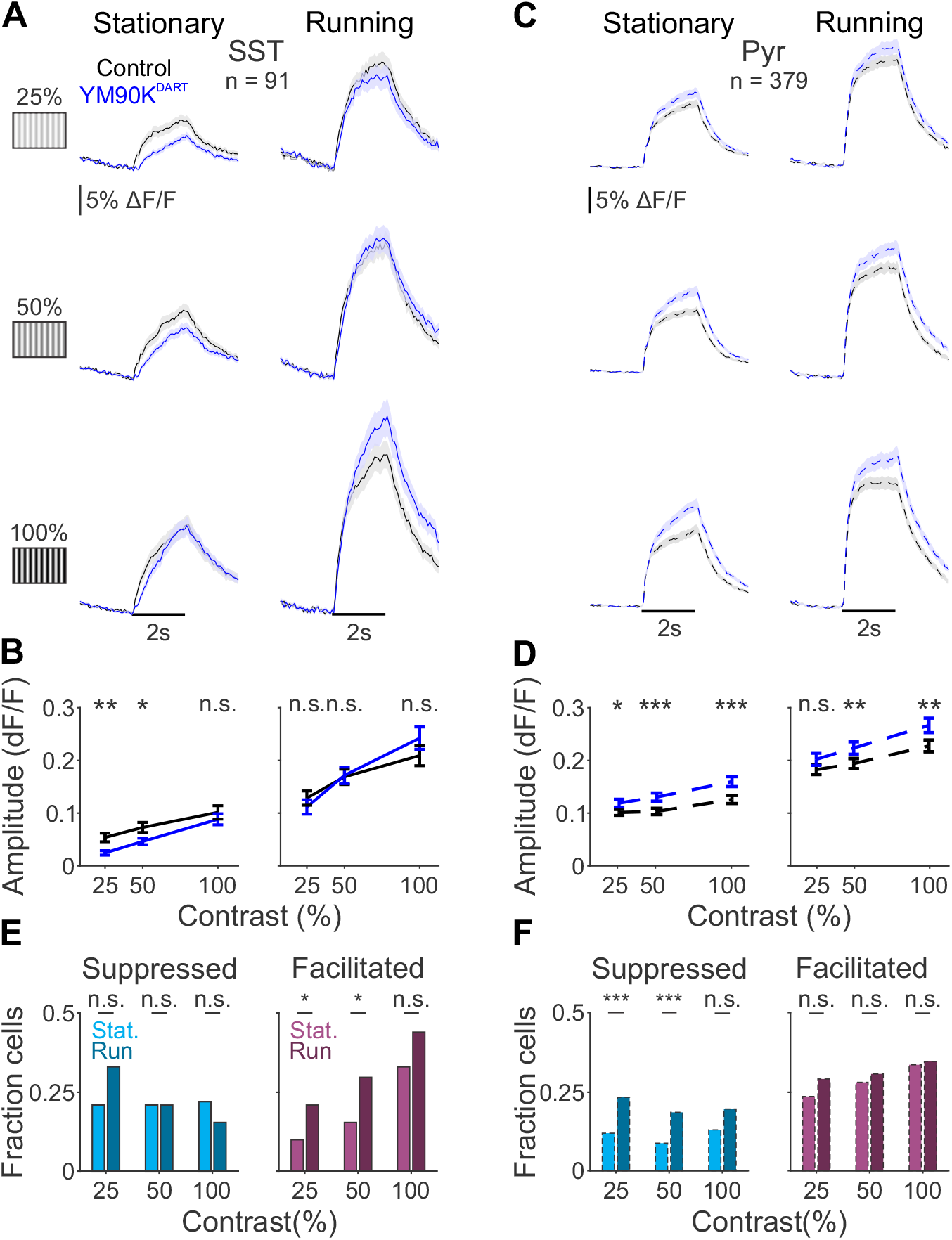
The effect of blocking AMPARs on SST cells depends on behavioral state. (A) Grand average time courses for SST cells before (black) and after (blue) YM90K^DART^ during stationary (left) or running (right) epochs, at each contrast. All cells are matched across behavioral states and contrasts. Shaded error represents SEM across cells. (B) Mean response during stimulus period, for SST cells during stationary (left) or running (right) epochs, at each contrast. Error is SEM across cells. (C-D) Same as (A-B), for pyramidal cells. (E) Fraction of SST cells suppressed (left, cyan) or facilitated (right, magenta) by more than 1 std of their control response during stationary (light) or running (dark) epochs. (F) Same as E, for pyramidal cells. n.s.-not significant; * p < 0.05; ** p < 0.01; *** p< 0.001. See also **Figure S5**.

The effects on the average responses are due to an increase in the fraction of SST cells facilitated by YM90K^DART^ when the mice are running (chi-square with Bonferroni correction for stationary vs. running, 25% contrast, p = 0.040; 50% contrast, p = 0.021; 100% contrast p = 0.128; **Figure 5E**), without a significant change in the fraction of cells suppressed (p > 0.05 for all contrasts). This variation in the effects of YM90K^DART^ on SST cells, with some being facilitated while others are suppressed, is consistent with our observation that SST cells are heterogenous in their contributions to stabilizing the network.

Arousal has also been linked to network changes in visual cortex activity, and is considered to be mechanistically distinct from the effects of locomotion^43–45^. To determine the impact of arousal on the network stabilizing role of SST cells, we segregated stationary trials according to pupil diameter^46,47^. For each mouse, we measured pupil size during stationary epochs across both experimental days and performed a median split on the trials to assign them to large and small pupil categories (**Figure S5A**). We confirmed that the average pupil diameter is significantly greater in the large pupil trials, (paired t-test p < 0.001, **Figure S5B**), and is similar to the size during locomotion (paired t-test, p = 0.079). To directly compare the same cells in each arousal state, we examined the subset of SST and putative pyramidal cells which could be measured at their preferred direction, in both small and large pupil conditions, for all contrasts, and during both imaging sessions (n = 10 mice; 107 SST cells and 468 pyramidal cells).

Arousal slightly, but significantly, facilitates responses of SST (three-way ANOVA, main effect for pupil size, p = 0.009) and pyramidal (three-way ANOVA, main effect for pupil size, p < 0.001) cells. As with locomotion, we find that arousal alters the effect of YM90K^DART^ on SST cells (three-way ANOVA, YM90K^DART^ x pupil size interaction p < 0.001). Specifically, SST cells are suppressed by YM90K^DART^ during low-arousal trials (small pupil trials: two-way ANOVA, main effect of YM90K^DART^, p < 0.001; **Figure S5C-D**), but not during high arousal trials (large pupil trials: p = 0.150). This is consistent with the arousal-dependent effects of YM90K^DART^ on pyramidal cells (three-way ANOVA, YM90K^DART^ x pupil size interaction p < 0.001). Pyramidal cells are disinhibited during low arousal (small pupil trials: two-way ANOVA, main effect of YM90K^DART^, p = 0.011; **Figure S5E-F**) and even more so during high arousal (large pupil trials: p < 0.001). These results suggest that recruitment of SST cells into the ISN is enhanced not only by stimulus strength, but also active states such as locomotion and arousal.

### Stimulus strength and behavioral state recruit SST cells into the ISN through distinct effects on the network

Our experimental data is in broad agreement with the predictions of the theoretical model, indicating that SST cells are increasingly needed for stabilization as network activity increases. To investigate how changing stimulus strength and behavioral state may act to engage SST cells into network stabilization, we returned to our modeling framework and fit our model weights to the neural responses in control and YM90K^DART^. In our model, YM90K^DART^ solely affects *W*_*SE*_ (since we assume that the direct sensory input to S cells (*I*_*S*_) is negligible^27^), and is modelled as a fractional change of this weight, (1 − *x*)*W*_*SE*_. We set *x* to 0.5 (**Figure S6**) as this is a conservative estimate of the efficacy of YM90K^DART^ based on our *in vitro* recordings. We modelled changes in contrast by changing the external inputs, *J*_*E*_ and *J*_*S*_, while holding all weights within the network constant, as we do not anticipate stimulus-dependent changes to synaptic connectivity. To model changes in behavioral state, we allowed both external inputs and weights to vary, capturing locomotion-dependent effects both on the strength of feedforward excitation and on synaptic connectivity within the network^34,48^.

We find that when all weights are allowed to vary between stationary and running conditions, the model can quantitatively fit the data very well (“Full model,” **Figures 6A** and **S6** and **Table S1**). We next investigated whether a more circumscribed set of flexible parameters could also capture the data. Multiple studies have highlighted VIP cells in regulating SST cell activity^29,30^, which has been proposed as a mechanism of locomotion modulation^31^. Therefore, we tested a model in which only the external inputs and the gain of V cells were permitted to change between behavioral states. To do so, we fit a gain term *g* that was applied to *W*_*SVS*_ and *W*_*SVE*_, with all other weights fixed. This model is also quite successful in fitting the data (“VIP model,” **Figure 6A,C** and **Table 1**) and produces a lower Akaike information criterion (AIC) value than the Full model (**Figure 6B**). Finally, to confirm that the change in V parameters are necessary, we compared this to a model where only external inputs could vary between states (“Input model”). This results in a higher cost and AIC value than the VIP model (**Figures 6A-B** and **S6** and **Table S2**). Thus, we focused on the VIP model since it yields the best fit according to AIC.

**Table 1.** Best fit parameters for V1 network model, when weights do not depend on state, except through changes in VIP gain. Optimal weights identified by our fitting procedure (**STAR Methods**) for effective connectivity within the V1 network (*W*_*EE*_ − *W*_*EPE*_ through *W*_*SVS*_) and external inputs (*J*_E_ and *J*_*S*_). Abbreviations as in **Figure S1**. External inputs vary with stimulus contrast and network weights are constant, except through changes in VIP gain (*g* = 2.387) during running. See also **Tables S1-2**.

| *Parameters* | *Stationary* |  |  | *Running* |  |  |
| --- | --- | --- | --- | --- | --- | --- |
|  | 25% | 50% | 100% | 25% | 50% | 100% |
| $W_{EE} - W_{EPE}$ | 0.996 | | | | | |
| $0W_{ES} - W_{EPS}$ | 0.257 | | | | | |
| $W_{SE}$ | 3.601 | | | | | |
| $W_{SVE}$ | -4.189 | | | -4.189 $g = -9.999$ | | |
| $W_{SVS}$ | 0.177 | | | 0.177 $g = 0.422$ | | |
| $J_E$ | 0.176 | 0.179 | 0.188 | 0.211 | 0.215 | 0.218 |
| $J_S$ | -0.409 | -0.409 | -0.518 | -0.032 | -0.032 | -0.067 |

**Figure 6.**
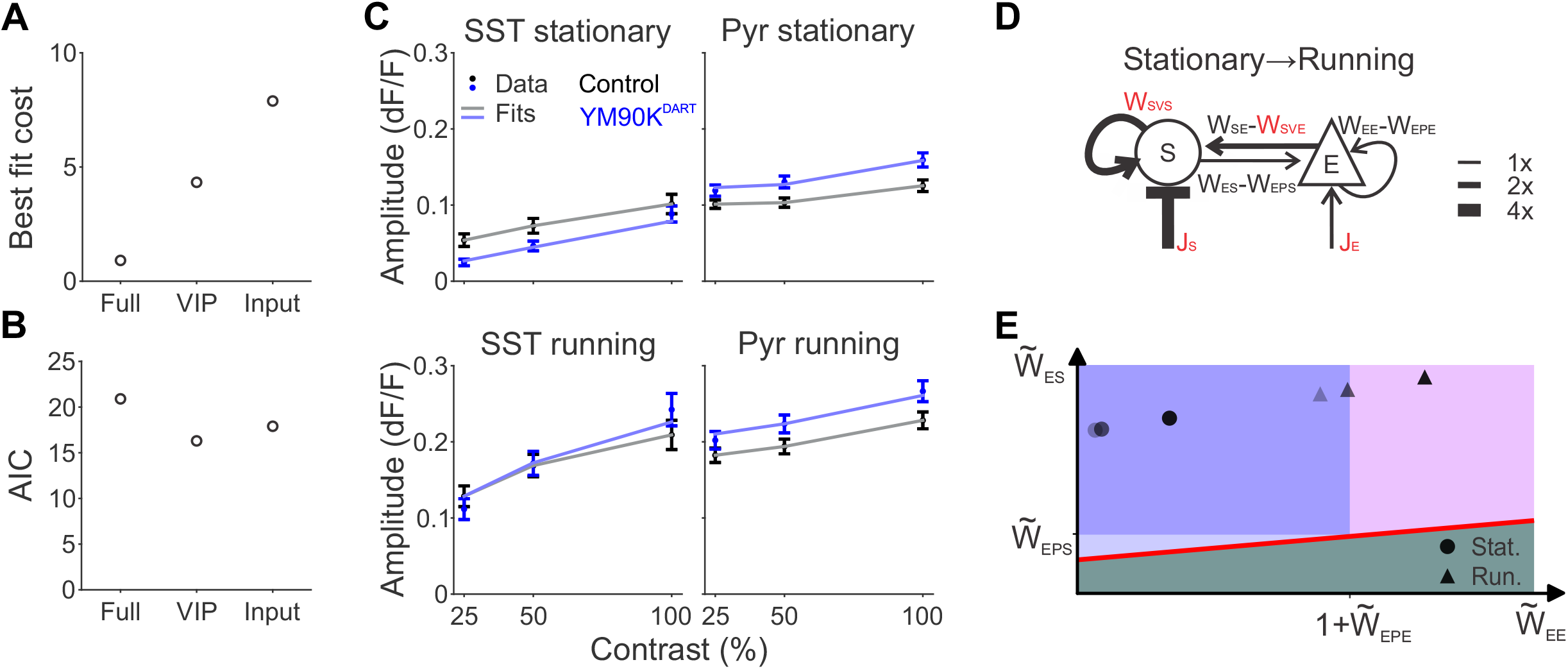
Paradoxical effects indicate the necessity of SST cells for network stabilization. Cost of the best fit for each of the three models. (B) Akaike information criterion (AIC) values for each of the three models. (C) Empirical (dark data points, mean +/-SEM from **Figure 5B,D**) and simulated (light lines) responses of SST (left) and pyramidal (right) cells to increasing contrast, in stationary (top) or locomotion (bottom) states in control (gray) and after YM90K^DART^ (light blue). (D) Schematic of changes to weights to fit changes from stationary to running. Line thickness is proportional to weight change. (E) Position of model best fit parameters at each contrast (shading) and behavioral state (circles = stationary, triangles = running) in the phase space from **Figure 1**. Instability line (red) corresponds to the high contrast, running condition. See also **Figure S6**.

The contrast-dependent effects of YM90K^DART^ on pyramidal and SST cells within each state are captured by changes in the net inputs *J*_*E*_ and *J*_*S*_ (**Figure 6C** and **Table 1**). *J*_*E*_ is positive and increases with contrast, consistent with increasing feedforward input. In comparison, *J*_*S*_ is always negative, reflecting increased inhibition to S cells from V cells, as our model includes no direct sensory input to S (**Figure S1**). Additionally, *J*_*S*_ decreases with contrast, consistent with increased input to V cells with increasing stimulus strength.

In the transition from the stationary to locomotion states, the increased gain to V cells (by a factor *g*) increases *W*_*SVS*_ such that the net recurrent effect of S cells through V cells is more excitatory (**Figure 6D** and **Table 1**). Meanwhile, *W*_*SVE*_ becomes more negative, such that *W*_*SE*_ - *W*_*SVE*_, the net excitation from E to S cells, also increases. Finally, the external input to E cells (*J*_*E*_) is elevated during locomotion. Thus, despite the decrease in *J*_*S*_, the net effects combine to increase recurrent excitation of S cells alongside higher activity of E cells during locomotion.

Plotting the network with our fit parameters in the 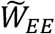 vs. 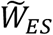 space defined in **Figure 1** allows us to gain insight into how the empirical network moves as a function of input strength and behavioral state (**Figure 6E**). We find that that the network is in R_i_ (i.e., the region in which S cells are not required for stability) in stationary conditions. As contrast increases, the network moves toward the boundary between R_i_ and R_ii_; that is, the effective recurrent excitation approaches the value at which it can no longer be stabilized by PV cells alone. Running shifts the network closer still to the R_i_-R_ii_ boundary, and when running coincides with high contrast the network crosses the border into R_ii_. Thus, high contrast stimuli during active epochs produce a network state in which P cells are insufficient to balance the effective recurrent excitation of pyramidal cells, and S cells are required to prevent network instability. We also find that the strength of inhibition of S cells onto pyramidal cells exceeds the disinhibition they provide through P cells.

Thus, the model supports our interpretation that YM90K^DART^ reveals the conditions under which SST cells contribute to the ISN, and that distinct mechanisms underlie the recruitment of SST cells to the ISN with increasing contrast or locomotion. Specifically, while contrast alters network activity directly through increased feedforward input, running additionally changes the local connectivity weights within V1, potentially via its action on VIP cells.

## Discussion

Inhibition stabilized networks^4–7^ are proposed to enable sensory cortex to normalize responses across a broad range of contexts. The data presented here provide insight into how the diverse cell types in the visual cortex circuit enable this flexibility. Employing cell-type specific pharmacology to reduce excitatory input to SST cells, and a novel theoretical framework for understanding this manipulation, we reveal that SST cells are required for network stabilization in mouse V1 under select conditions of high sensory drive and active behavioral states. This work provides a concrete example for how different cell types play complementary roles in regulating sensory processing across stimulus and behavioral contexts.

The major innovation that enabled these experiments is the ability to selectively block synaptic excitation onto SST cells by using the AMPAR antagonist YM90K^DART^. This offers several advantages over more typical methods for manipulating neuronal activity to probe ISNs, such as optogenetic activation of somato-dendritic conductances^8–10,13,14^. First, YM90K^DART^ allows us to directly manipulate a circuit feature that is critical to recruiting recurrent inhibition, namely the recurrent excitation from pyramidal to SST cells (*W*_*SE*_). Second, unlike optogenetic activation of conductances, the efficacy of YM90K^DART^ does not depend on neuronal excitability (e.g., distance from threshold and input resistance), which is impacted by both changing sensory input and behavioral state. Thus, YM90K^DART^ enables a more straightforward interpretation of the apparent decrease in efficacy of our manipulation with increasing stimulus strength and arousal.

When mice are quiescent and visual stimuli are weak, YM90K^DART^ reduces SST cells’ responses, while moderately disinhibiting responses of putative pyramidal cells. This intuitively expected effect is consistent with past work highlighting the role of PV cells in network stabilization during both spontaneous activity and sensory integration^5,10,13,14,16^. However, our model and others’^25^ argue that there is a limit to the strength of recurrent excitation that the PV cells can stabilize, and that past this point SST cells are also needed for network stabilization. Indeed, when visual stimuli are strong, we find that decreasing excitation onto SST cells elicits stronger disinhibition of excitatory cells, and a paradoxical facilitation of an increasing number of SST cells.

Our model recapitulates this contrast-dependent effects of YM90K^DART^ on both pyramidal and SST cells solely through changes in the sensory inputs to these cell types (*J*_*E*_ and *J*_*S*_). In our model, contrast-dependent effects arise due to a non-linearity of the input-output transfer function, but other non-linearities, such as those introduced by short-term plasticity, could also play a role.

Notably, this facilitation of SST cells could not occur if YM90K^DART^ blocked all excitatory input to SST cells. The remaining excitatory input may be mediated by a subset of unblocked AMPARs. Indeed, our *in vitro* electrophysiology recordings demonstrate substantial, but not complete, reduction of AMPAR-mediated excitation on SST cells, and suggest that a subset of synapses may remain intact. In addition, excitatory input to SST cells is facilitating and thus may be more effectively recruited by the higher frequency firing evoked with increasing stimulus strength. Alternatively, non-AMPARs such as NMDARs and metabotropic glutamate receptors are both expressed on SST cells, and may also provide a source of continued excitatory drive in the presence of YM90K^DART^.

Our finding that the effect of YM90K^DART^ depends on the correlation of each SST cell’s activity with the local network supports our conclusion that we are revealing their engagement in the ISN. In an ISN, only those cells that are strongly coupled to the network should be facilitated by the disinhibitory effects of YM90K^DART^, whereas weakly coupled neurons undergo net suppression. Surprisingly, we also found that the network coupling of SST cells predicted the strength of their visual stimulus responses in the control condition, where weakly correlated cells were more robustly driven. One possibility is that weakly correlated SST cells receive less recurrent excitation and are more strongly driven by long-range inputs. Given that we presented full-field gratings, SST cells receiving long range inputs may be more effectively driven, and less surround suppressed, by these stimuli. Future experiments taking advantage of genetic access to molecularly distinct subtypes of SST cells will be helpful in understanding the origins of this heterogeneity. Nonetheless, these results suggest that the transition from a purely PV stabilized network to a SST stabilized network may be a gradual process with the progressively stronger recruitment of SST cells into the ISN.

We also find that behavioral state critically controls the recruitment of SST cells into the ISN. Locomotion dramatically increases stimulus responses of all major cell types in the V1 circuit ^32,34,42^, and models suggest that visual stimulation coupled with locomotion creates the conditions for SST recruitment to the ISN^25,26^. Indeed, when mice are running, we find limited suppression of SST cells by YM90K^DART^ even with low contrast stimuli. When the mice run during high contrast stimuli, we observe clear paradoxical facilitation. Given that increasing stimulus strength, locomotion and arousal all drive stronger visually-evoked activity in the pyramidal cells, it is possible that all of these conditions increase engagement of SST cells through the same mechanisms. However, arousal and locomotion are also associated with neuromodulation of V1, including by cholinergic and noradrenergic inputs^31,50,51^. By altering cells’ excitability and synaptic output, neuromodulation could effectively change the connectivity weights in the V1 circuit^34^, pushing the network into a region in which SST cells are required for stabilization. Consistent with the literature, our model suggests that this may occur through modulation of VIP cells^31^.

While the finding that SST cells are recruited into the ISN during presentation of strong sensory stimuli and during states of behavioral arousal is likely to broadly generalize, the conditions that determine the transition between states will depend on the specific architecture of each cortical area. As the density of different cell types and their connectivity varies across the cortex, so will the boundaries between ISN regions. An important question for future inquiry is to understand how the transition from a purely PV-stabilized to a PV-and-SST-stabilized network impacts sensory processing. One possibility is that this transition has little effect on the input-output function of the excitatory population, and simply enables the network to maintain stability across a broader range of contexts. Alternatively, the transition to reliance upon dendrite-targeting SST cells may alter the dynamics of synaptic integration^49–51^ and plasticity, and may be finely tuned within each cortical area. Synapse and cell-type specific pharmacology coupled with our modeling framework promise to reveal how each node in the cortical circuit supports sensory processing across a broad range of environmental and behavioral contexts.

## Acknowledgements

We thank Wenjuan Kong, Lou Campillo, Tingwei Hu, Lindsey Wilson, T.J. Wagner, Gloria Kim, Dr. Sasha Burwell, and Megan Stone for assistance with husbandry, surgeries, retinotopic mapping and histology. We thank Dr. Ashley Wilson for initial characterization of DART delivery and expression in V1, Dr. Mark Histed for the gift of a virus for expression of GCaMP8s, Dr. Kevin Franks for the use of his epifluorescence microscope, and Dr. Yuansi Chen for advice on statistical approaches. We thank Dr. Court Hull for comments on the manuscript, and Dr. Rich Mooney and members of the Hull and Glickfeld labs for insight throughout the project. This work was supported by grants from the National Institutes of Health (R01-EY031716 to L.L.G., F32-EY034013 to C.M.C., F31-EY031941 to J.Y.L. and RF1-MH117055, DP2-MH1194025, R01-NS107472, and R61-DA051530 to M.R.T.), the American Heart Association (to S.S.X.L) and the Holland-Trice Foundation (to L.L.G.).

## Author Contributions

Conceptualization: C.M.C, N.B., M.R.T., and L.L.G.; Methodology: T.H., B.C.S. and S.S.X.L.; Investigation and Data curation: C.M.C., J.Y.L.; Formal analysis: C.M.C., Y.P., D.S.A, J.Y.L.; Writing-Original Draft: C.M.C. and L.L.G.; Writing-Review and Editing: C.M.C, J.Y.L., N.B., M.R.T., and L.L.G.; Visualization: C.M.C, Y.P. and L.L.G.; Supervision: C.M.C., N.B., M.R.T., and L.L.G.; Funding Acquisition: C.M.C, M.R.T., and L.L.G.

## STAR Methods

### Key Resources Table

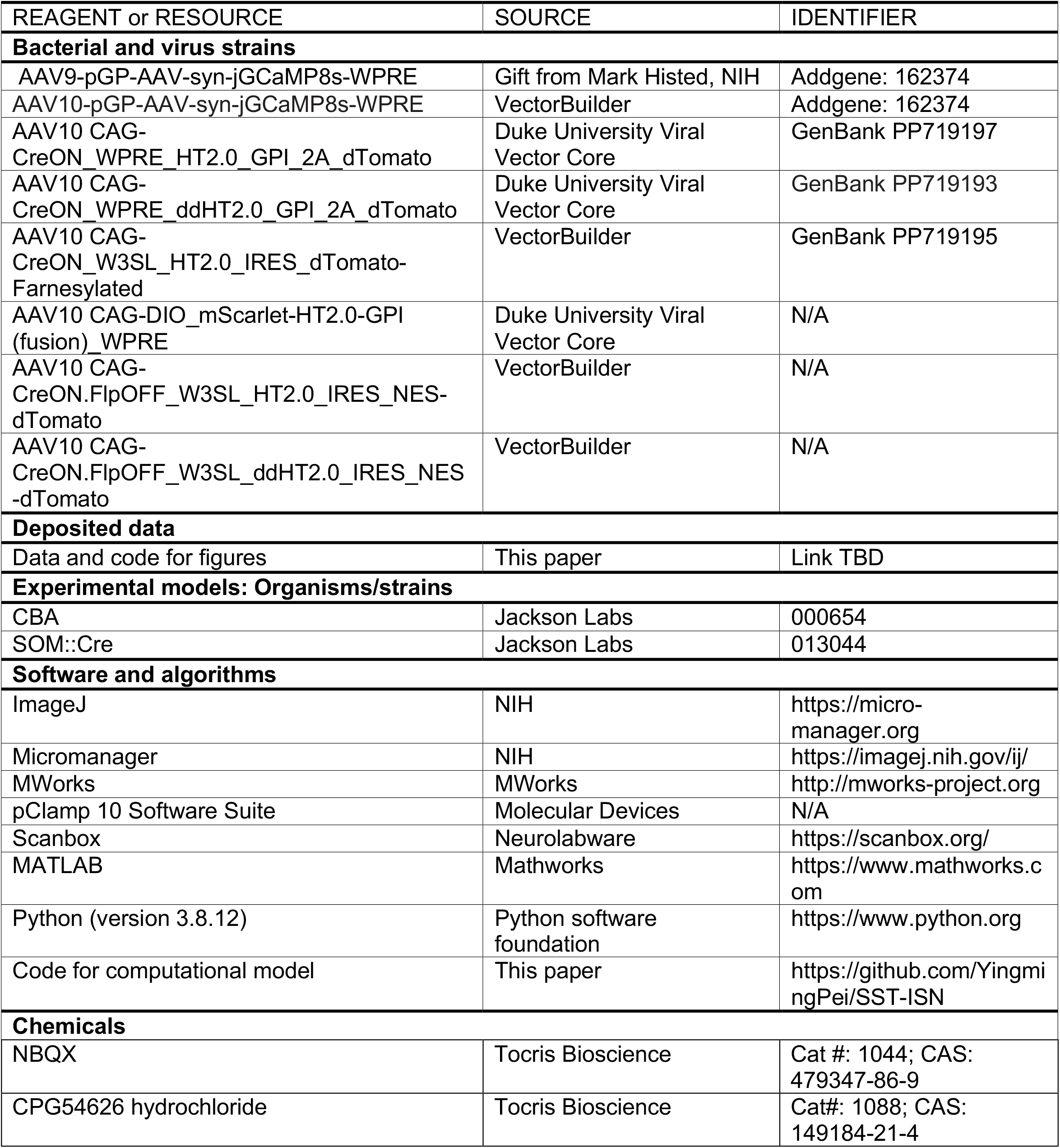
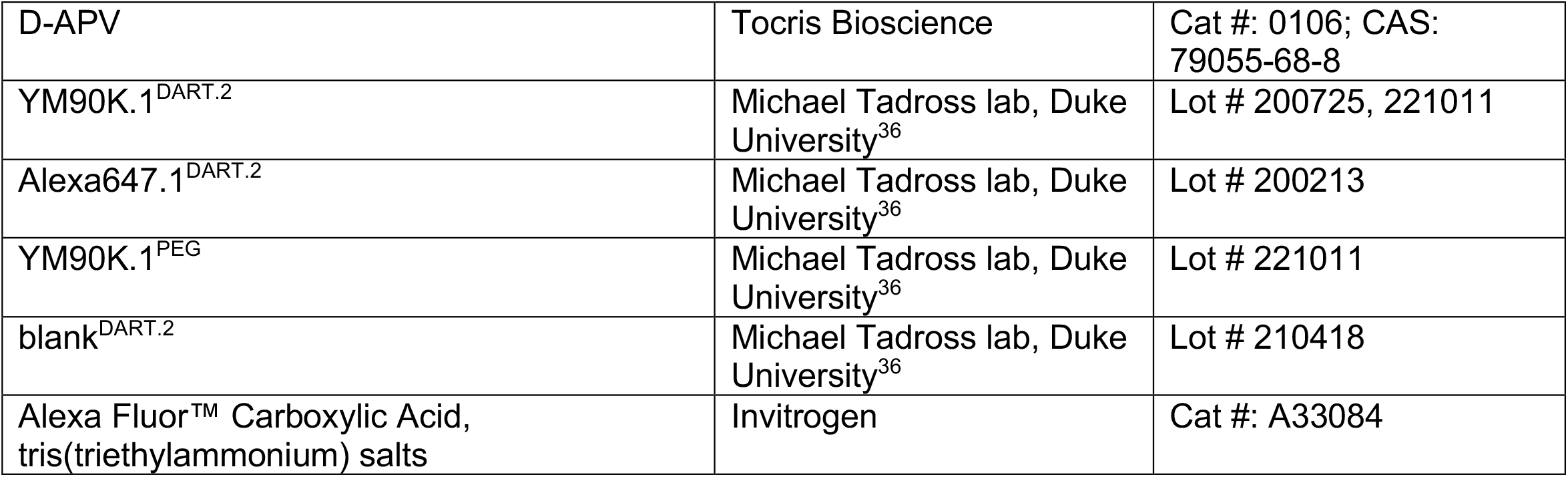

## RESOURCE AVAILABILITY

### Lead contact

Further information and requests for resources and reagents should be directed to Lindsey Glickfeld.

### Materials availability

No new reagents were generated as a result of this study.

### Data and code availability

All two-photon imaging data included in the manuscript figures is available on Figshare. A link is provided in the *Key resources table*.

All original code needed to generate the manuscript figures is available on Figshare. A link is provided in the *Key resources table*. The complete code for the computational model is available on Github. A link is provided in the *Key resources table*.

Any additional information required to reanalyze the data reported in this paper is available from the lead contact upon request.

## EXPERIMENTAL MODEL AND SUBJECT DETAILS

### Animals

All procedures conformed to standards set forth by the National Institutes of Health Guide for the Care and Use of Laboratory Animals, and were approved by the Duke University’s Animal Care and Use Committee. Mice were housed on a normal 12:12 light-dark cycle. Two-photon calcium imaging data in this study were collected from 13 mice (8 female). Of these, 8 mice were used only in YM90K^DART^ experiments, 3 mice were used only in YM90K^PEG^ experiments, and 3 mice were shared. Imaging experiments were conducted at 21-38 weeks of age (mean 31 weeks), except for one mouse imaged at 11 weeks. Headpost, cranial window, and cannula implantation were performed no earlier than 7 weeks, with viral injection a minimum of 3 weeks after. Electrophysiology data were collected from 22 mice (13 female). Electrophysiology experiments were conducted at 5-9 weeks of age. Viral injections for electrophysiology experiments were performed no earlier than 3 weeks of age. All mice for two-photon experiments were either offspring of CBA mice (Jackson Labs, #000654) crossed with SOM::Cre mice (Jackson Labs, #013044), or offspring of SOM::Cre mice crossed with PV::Flp (Jackson Labs, #022730). Mice used for electrophysiology experiments were of these two genotypes, or offspring of SOM::Cre mice crossed with R26R-EYFP mice (Jackson Labs, #006148) or crossed with Ai148 mice (Jackson Labs, #030328).

## METHOD DETAILS

### Surgical Procedures

#### Viruses

Due to the evolving nature of the novel DART reagents^35,36^, we used several constructs for HaloTag protein (HTP) and GCaMP expression over the course of data collection. We have found these to be functionally equivalent. In the methods, viruses are referenced by their identifiers in the following table:

| *Virus* | *Titer (GC/mL)* | *Identifier* |
| --- | --- | --- |
| AAV9-pGP-AAV-syn-jGCaMP8s-WPRE | 4.03x13 | GC1 |
| AAV10-pGP-AAV-syn-jGCaMP8s-WPRE | 6.00x13 | GC2 |
| AAV10 CAG-CreON_WPRE_HT2.0_GPI_2A_dTomato | 2.80x13 | HTP1 |
| AAV10 CAG-CreON_WPRE_ddHT2.0_GPI_2A_dTomato | 3.50x13 | ddHTP1 |
| AAV10 CAG-CreON_W3SL_HT2.0_IRES_dTom-Farnesylated | 9.00x12 | HTP2 |
| AAV10 CAG-DIO_mScarlet-HT2.0-GPI (fusion)_WPRE | 3.07x13 | HTP3 |
| AAV10 CAG-CreON.FlpOFF_W3SL_HT2.0_IRES_NES-dTom | 2.96x13 | HTP4 |
| AAV10 CAG-CreON.FlpOFF_W3SL_ddHT2.0_IRES_NES-dTom | 1.86x13 | ddHTP2 |

### Intracranial viral injections for electrophysiology

Burrhole injections of viral constructs (HTP1-4, ddHTP1-2) were used to express HTP for slice electrophysiology experiments. Mice were anesthetized with isoflurane (1.2-2% in 100% O2) and positioned in a stereotax (Kopf Instruments). Meloxicam (5 mg/kg) was administered subcutaneously and bupivacaine (5 mg/kg) was administered locally prior to incision. After the skull was exposed, a small hole was drilled +/-2.6 mm lateral from lambda and directly anterior to the lambdoid suture targeting the posterior and medial aspect of the primary visual cortex (V1). Injection micropipettes were pulled from glass capillary tubes (1B100F-4, World Precision Instruments) and backfilled with virus and then mineral oil and mounted on a Hamilton syringe. The pipette was lowered into the brain and 100-200 nL of virus was pressure injected at 10-40 nL/min using an UltraMicroPump (World Precisions Instruments) 200-250 µm below the surface. We waited 2-3 weeks for viral expression.

### Cisterna magna infusion for electrophysiology

For electrophysiology experiments with systemic DART, we introduced 2 μL YM90K^DART^ (3 mM) and Alexa647^DART^ (0.3 mM) to the cerebrospinal fluid acutely through injection to the cisterna magna. Meloxicam (2.5 mg/kg, s.c.) was administered at the start of the surgery. Animals were anesthetized with isoflurane (1.2-2% in 100% O2). An incision was made at the midline at base of the skull and muscle was displaced by blunt dissection until the membrane of the cisterna magna was accessible. The cisterna magna was located by visual identification. A small puncture was made in the cisterna magna membrane, and 2-5µL of the DART mixture was injected via a 30G needle mounted on a Hamilton syringe. The muscle was replaced and the skin was sutured. Buprenorphine (0.05 mg/kg, s.c.) was delivered upon recovery from anesthesia. Slices for electrophysiology were prepared 2.5-3 h after the cisterna magna injection.

### Cranial window implant

Animals were implanted with a titanium headpost and 3-5 mm cranial window. Dexamethasone (3.2 mg/kg, s.c.) and Meloxicam (2.5 mg/kg, s.c.) were administered at least 2 h before surgery. Animals were anesthetized with ketamine (200 mg/kg, i.p.), xylazine (30 mg/kg, i.p.) and isoflurane (1.2-2% in 100% O2). A midline incision was made to expose the skull, and muscle and membranous tissue were scraped away from the exposed bone. A guide cannula (F11552, P1 Technologies) with a complementary dummy cannula (F11372, P1 Technologies) was directed to the right lateral ventricle using the following coordinates from bregma: 1.10 mm lateral, 0.20 mm posterior, 2.30 mm from the skull surface. The cannula was secured to the skull with C&B Metabond (Parkell). Within the same surgery, a titanium headpost was secured using cyanoacrylate glue and Metabond, and a 3-5 mm craniotomy was made over the left hemisphere (center: 2.8 mm lateral, 0.5 mm anterior to lambda) allowing implantation of a glass window (a 5-8 mm coverslip bonded to two 3-5 mm coverslips (Warner no. 1) with refractive index-matched adhesive (Norland no. 71)) using Metabond. Buprenorphine (0.05 mg/kg) and cefazolin (50 mg/kg) were delivered s.c. every 12 h for 48 h following surgery. Mice were allowed to recover from surgery for a minimum of 7 d before subsequent procedures.

### Retinotopic mapping

Following at least 7 d recovery from the headpost implantation surgery, mice were gradually habituated to head restraint. After habituation, mice underwent retinotopic mapping using intrinsic autofluorescence imaging to locate V1 for viral injection. The brain was illuminated with white light (Lumen Dynamics, X-Cite 120) with a 472 ± 30 nm band pass filter (Edmund Optics), and emitted light was measured through a green and red filter (500 nm longpass). Drifting gratings were presented on a monitor positioned at 45° relative to the body axis, and stimuli were shown at 3 positions (Elevation: -10 deg, Azimuth: -30, 0, and 30 deg, 45° diameter with a gaussian mask, drifting at 2 Hz, 10 s duration, 10 s inter-trial interval (ITI)) to activate locations in the contralateral visual field. Images were collected using a CCD camera (Rolera EMC-2, QImaging) at 2 Hz through a 5x air immersion objective (0.14 numerical aperture (NA), Mitutoyo), using Micromanager acquisition software (NIH). Images were analyzed in ImageJ (NIH) to measure changes in fluorescence (dF/F; with F being the average of all frames). Injections were targeted to the region of V1 driven by the center position.

### Viral injections for two-photon imaging

The mice used for two-photon imaging underwent an additional surgery for viral injection. Dexamethasone (3.2 mg/kg, s.c.) was administered at least 2 h before surgery. After anesthesia with isoflurane (1.25–2% in 100% O2), the cranial window was removed. HaloTag virus (HTP 2-4) mixed with GCaMP8s (GC 1-2) in a 1:1 ratio was injected via a glass micropipette mounted on a Hamilton syringe. Two hundred to three hundred nanoliters of virus were injected at 170-230 µM below the pia (30 nL/min); the pipette was left in the brain for an additional 3 min to allow the virus to infuse into the tissue. Following injection, a new coverslip was sealed in place with Metabond. We then waiting a minimum of two weeks for viral expression to mature before performing two-photon experiments.

## Experimental Procedures

### In vitro slice preparation

Mice were deeply anesthetized with isoflurane, the brain was removed and then transferred to oxygenated (95% O_2_ and 5% CO_2_), ice-cold artificial cerebrospinal fluid (ACSF, in mM: 126 NaCl, 2.5 KCl, 26 NaHCO_3_, 1.25 NaH_2_PO_4_, 20 glucose, 2 CaCl_2_, 1.3 MgCl_2_). Coronal brain slices (300 µm thickness) were prepared using a vibrating microtome (VT1200S, Leica) and transferred to a holding solution (at 34º C) for 12 min, and then transferred to storage solution for 30 min before being brought to room temperature. The holding solution contained (in mM): 92 NaCl, 2.5 KCl, 1.25 NaH_2_PO_4_, 30 NaHCO_3_, 20 HEPES, 25 glucose, 2 thiourea, 5 Na-ascorbate, 3 Na-pyruvate, 2 CaCl_2_, 2 MgSO_4_. The storage solution contained (in mM): 93 NMDG, 2.5 KCl, 1.2 NaH_2_PO_4_, 30 NaHCO_3_, 20 HEPES, 25 glucose, 2 thiourea, 5 Na-ascorbate, 3 Na-pyruvate, 0.5 CaCl_2_, 10 MgSO_4_. For DART incubation (0.5-4 h) we used the same holding solution, with the addition of 1 µM YM90K^DART.2^ and 0.1 µM Alexa647^DART.2^. Additional controls used this holding solution with 1 µM blank^DART.2^ or 1 µM YM90K.1^PEG^. Micropipettes pulled from borosilicate glass (1B150F-4, World Precision Instruments) were filled with internal solution containing (in mM): 142 K-gluconate, 3 KCl, 10 HEPES, 0.5 EGTA, 5 phosphocreatine-tris, 5 phosphocreatine-Na2, 3 Mg-ATP, 0.5 GTP. Recording pipettes had resistances of 3-10 MΩ.

### In vitro slice recordings

Recordings occurred between 1.5 and 5 h after the animal was sacrificed. Brain slices were transferred to a recording chamber and maintained at 34º C in oxygenated ACSF (containing, in mM: 136 NaCl, 2.5 KCl, 26 NaHCO_3_, 1.25 NaH_2_PO_4_, 20 glucose, 2 CaCl_2_, 1.3 MgCl_2_, bubbled with 95% O_2_ and 5% CO_2_) perfused at 2 mL/min. Electrophysiological recordings were restricted to layer 2/3 and V1 was identified by visualization of fluorescence expression at the viral injection site. Neural signals were recorded using a MultiClamp 700B and digitized with a Digidata 1550 (Axon Instruments) with a 20 kHz sample rate. Data acquisition and stimulus presentation was controlled using the Clampex software package (pClamp 10.5, Axon Instruments).

In voltage-clamp recordings, series resistance was monitored using -5 mV steps preceding each trial. Only cells that had < 30 MΩ series resistance were included in analysis. Spontaneous EPSCs (**Figure 2**) were recorded from SST cells, identified by dTomato expression, with cells held at a membrane potential of -85 mV to isolate excitatory events. Following a minimum of 2.5 min in normal ACSF, we washed on NBQX (10 µM, TOCRIS Bioscience) and allowed 2.5 min for NBQX to saturate the slice before collecting data in this condition. To compare EPSC amplitude in SST and putative pyramidal cells (**Figure S2**), we patched nearby pairs (< 50 µm distance) and identified cells based on dTomato expression and somatodendritic morphology. EPSCs were evoked by electrical stimulation (150-250 µA; 100 µs duration) with a steel monopolar electrode placed in layer 2/3 in between the recorded cells (∼100 µm distance from each cell to electrode). Stimulation location and intensity were adjusted prior to data collection to minimize polysynaptic activation (assessed with online observation of EPSCs). Based on our previous data silencing local action potentials with muscimol, we considered monosynaptic responses to be short-latency (< 5 ms) EPSCs^52^. All recordings were performed in ACSF containing MCPG (0.4 mM), CGP54626 (1 µM), and APV (30 µM) to block mGluRs, GABA_B_Rs and NMDARs, respectively. In a subset of these experiments, DART reagents (300 nM YM90K^DART^ and 100 nM Alexa647^DART^) were applied acutely (**Figure S2A-B**); in the remainder, DART reagents were infused via the cisterna magna (**Figure S2C-D**). All data are the average of a minimum of 10 trials.

### Intracerebroventricular (ICV) infusion

2 µL YM90K^DART^ (3 mM) was co-infused with Alexa647^DART^ (0.3 mM), while the non-binding YM90K^PEG^ (3 mM) was co-infused with Alexa647-COOH (0.3 mM). During infusion, mice were headfixed on a running wheel and the dummy cannula removed. An internal cannula (F11373, P1 Technologies) connected to a Hamilton syringe on an infusion pump was inserted into the guide cannula and secured in place. Compounds were delivered at 75-100 nL/min, followed by at 10-20 min waiting period before the internal cannula was removed. The dummy cannula was then reinserted and secured. For the mice used in both YM90K^DART^ and YM90K^PEG^ experiments, the YM90K^PEG^ infusion and two-photon data collection were always performed at least 48 h prior to the pre-YM90K^DART^ control session.

We visualized Alexa647^DART^ and Alexa647-COOH through the cranial window using widefield microscopy. The brain was illuminated with orange light via a 624 ± 40 nm band pass filter (Edmund Optics) through the cranial window and far-red fluorescence was collected through a 692 ± 40 nm band pass filter (Edmund Optics). Images were collected using a CCD camera (Rolera EMC-2, QImaging) through a 5X air-immersion objective (0.14 numerical aperture (NA), Mitutoyo) using Micromanager acquisition software (NIH).

### Two-photon imaging

Images were collected using a two-photon microscope controlled by Scanbox software (Neurolabware). A Mai Tai eHP DeepSee laser (Newport) was directed into a modulator (Conoptics) and raster scanned on the visual cortex using resonant galvanometers (8 kHz; Cambridge Technology) through a 16X (0.8 NA, Nikon) water-immersion lens at a frame rate of 15 Hz. Emitted photons were directed through a green (510 ± 42 nm band filter; Semrock) or red filter (607 ± 70 nm band filter; Semrock) onto GaAsP photomultipliers (H10770B-40, Hamamatsu). At the start of each experiment, we used an excitation wavelength of 1040 nm to visualize dTomato fluorescence, allowing identification of red SST cells. All functional imaging used an excitation wavelength of 920 nm. Data were collected at 175 – 250 µM below the cortical surface.

During imaging experiments, mice were head-fixed and allowed to freely run on a cylindrical treadmill. Running speed was monitored with a digital encoder (US Digital). Pupil position was monitored via scattered infrared light from two-photon imaging. Light was collected using a GENIE Nano CMOS camera (Teledyne Dalsa) using a long-pass filter (695 nm) at the imaging rate. For each mouse we performed a baseline imaging session prior to the ICV infusion, and performed a second imaging session 17-24 h later, finding the same plane as in the baseline session using the vasculature and HTP expression as fiduciary markers.

### Visual stimulus presentation

Visual stimuli were presented on a 144-Hz (Asus). The monitor was calibrated with an i1 Display Pro (X-rite) for mean luminance at 50 cd/m2 and positioned 21 cm from the eye. Stimuli were generated and displayed using MWorks (The MWorks Project).

At the beginning of each session, we performed a retinotopy (9 positions, 30 deg diameter gabor grating, 15 deg spacing in azimuth and elevation) to position the monitor such that the receptive fields of the imaged neurons were centered on the screen. During the experiment, full-field, sine-wave gratings (0.1 cycles per degree; 2 Hz) were randomly interleaved at 3 contrasts (25, 50 and 100%) drifting in 8 directions (45 deg increments) for 2 s. Stimuli alternated with a 4 s ITI of uniform mean luminance (60 cd/m^2^).

### Post-hoc histology

After recording, animals were anesthetized with an overdose of ketamine (50 mg/kg) and xylazine (5 mg/kg) and perfused with PBS followed by 4% PFA in PBS. Brains were dissected and incubated in 4% PFA overnight, rinsed 3x with PBS, then sliced in 70-100 µm sections and mounted on glass slides. Slides were mounted with Fluoromount G with DAPI (Invitrogen) and imaged using an epifluorescence microscope (Keynce BZ-X8100) to confirm overlap of viral expression (GCaMP: excitation-470 ± 40 nm., emission-525 ± 50 nm; dTomato: excitation-560 ± 40 nm., emission-630 ± 75 nm) and capture (Alexa647: excitation-605 ± 50 nm., emission-670 ± 50 nm), and appropriate placement of the cannula in the lateral ventricle.

## QUANTIFICATION AND STATISTICAL ANALYSIS

All analyses were performed using custom code written in MATLAB (Mathworks; for electrophysiology and imaging data) or Python (for computational modeling). N values refer to number of cells or mice. Sample sizes were not predetermined but were collected to be comparable to published literature for each type of experiment^29,53,55,57,59^. Our sample size differs depending on the specific comparison made, as we always used subsets of cells that could be compared across all conditions.

### Electrophysiology

#### Spontaneous EPSCs

Initial event detection was conducted using a template search in Clampfit (pClamp 10.5, Axon Instruments). Spurious events were rejected by visual inspection. Of the remaining events, we rejected those with an amplitude less than 15 pA or greater than 175 pA, or with a rise time greater than 1 nA/mS. These criteria were based on visual inspection of true events compared to noise. To determine the sEPSC rate, we counted the sEPSCs in each sweep and divided by the sweep length to find events per second, then calculated the average rate across sweeps in each condition. To find the sEPSC amplitude we calculated the mean of the event peak amplitude (from the template match) in each sweep, then calculated the mean across sweeps in each condition.

#### Analysis of evoked EPSCs

Amplitudes of EPSCs in response to electrical stimulation were quantified from the mean of the last 10 sweeps of each condition. Amplitudes were calculated as the average response in a 2 ms window around the peak of the response. Cells were excluded from analysis if the resistance changed by more than 20% over the course of the recording. The mean EPSC amplitude for each SST cell was compared to that of the putative pyramidal cell in the same pair to determine the SST:pyramidal EPSC ratio.

### Two-photon calcium imaging

#### Registration, segmentation, matching across sessions, and time course extraction

To adjust for x-y motion, we registered all frames from each imaging session to a stable reference image selected out of several 500-frame-average images, using Fourier domain subpixel 2D rigid body registration. For each experiment, we first segmented cells in the YM90K^DART^ session and then used this as a reference to find matching cells in the control session. Cells bodies were manually segmented, first using the dTomato fluorescence to identify HTP+ SST cells, then selecting all other visible cells from images of the average dF/F during stimulus presentation (where F is the average of 1 s preceding each stimulus) for each unique stimulus condition, a time-averaged image of F across the full stack, and a local correlation map (where the value of each pixel is scaled by its correlation with the neighboring 9 pixels). All segmented cells that were not identified based on dTomato fluorescence were labelled as HTP- and assumed to be putative pyramidal cells.

We then found matching cells in the control session. After registration, salient fiduciary marks (e.g. bright cells and thin vasculature) were used to align the image stack to the YM90K^DART^ session. Then, for each cell segmented in the YM90K^DART^ session we examined an approximately 24.5 × 34.5 µM FOV in the corresponding region of the stack from the control session to determine whether the matching cell was detectable. Matching cells were visually identified based on location and morphological similarity to the corresponding cell in the YM90K^DART^ session. Within the small FOV, we used either the dTomato fluorescence (for cells labelled as HTP+ SST in the YM90K^DART^ session), the local correlation map, the time-averaged F across the full stack, or the maximum dF/F projection to identify and manually segment cells in the control session matching those found in the YM90K^DART^ session. Fluorescence time courses were derived by averaging all pixels in a cell mask. To exclude signal from the neuropil, we first selected a three pixel shell around each neuron (excluding a three pixel boundary around the segmented neuron and the territory of neighboring neurons), then estimated the neuropil scaling factor by maximizing the skew of the resulting subtraction, and finally subtracted this component from each cell’s time course^54^.

#### Visual responses and cell inclusion

Visually-evoked responses were measured as the average dF/F in the 2 s stimulus period starting 3 frames (200 ms) after visual stimulus onset and ending 3 frames after stimulus offset to account for cortical response latency. Among cells that we could identify in both imaging sessions, we included cells that were visually responsive (demonstrated a statistically significant elevation in dF/F during the stimulus period for at least one stimulus condition as defined by a Bonferroni corrected paired t-test) in at least one of the sessions. We applied the additional criterion of excluding any cell that had a mean visually evoked response more than 3 standard deviations greater than the mean response of all cells in that imaging session. We then found the preferred direction of visual grating for each cell on each day by identifying the direction with the maximum dF/F response, and all analyses were performed on the subset of trials at that grating direction for each cell.

For analysis of locomotion and arousal, we used subsets of cells that were represented across all conditions. This required that each cell have trials at its preferred direction, for each contrast and state on both imaging sessions. When comparing stationary and locomotion conditions, this stringent inclusion criterion led to the loss of two animals from the YM90K^DART^ experiment and two from the YM90K^PEG^ experiment (these were not the same mice). When comparing small pupil and large pupil conditions, the inclusion criteria excluded a small number of cells, but did not result in the loss of any mice from the sample.

#### Normalized difference and fraction suppressed or facilitated

As a measure of the impact of YM90K^DART^ on each cell’s visual responses, we defined a normalized difference metric:

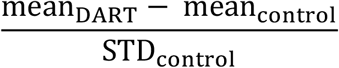

This normalization accounts for the difference in response magnitude across cells. The resulting metric is positive when a cell had a larger response in the YM90K^DART^ session and negative when the cell had a weaker response in the YM90K^DART^ session, compared to the control session. Cells were designated as “suppressed” if the normalized difference was <-1; that is, if the cell’s response in the DART session was more than one standard deviation below than that on the control day. Likewise, cells were designated as “facilitated” if the normalized difference was >1. The fraction of cells suppressed or facilitated was calculated by dividing the number of cells that met the above criteria by the total number of cells of that type.

For direct comparison of YM90K^DART^ and YM90K^PEG^ (**Figures S3-4)** we computed a modulation index for each neuron:

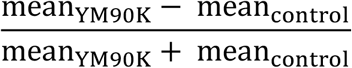

Cells that had a response <0 during either drug or control sessions were set to 0, so that values are restricted to be between -1 and 1.

#### SST-Pyr correlation

To separate SST cells into those strongly or weakly correlated with ongoing pyramidal activity, we first found the mean visual response of each SST cell, or the population of neighboring pyramidal cells, to every combination of contrast, direction, and behavioral state. This condition mean was then subtracted from the activity on each trial of that condition and used to calculate the Pearson correlation (using *corrcoef* in MATLAB) for each SST cell with the simultaneously imaged pyramidal population using only stationary trials on the control day. Cells with an R value greater than 0.5 were designated as “strongly correlated” and those with an R value less than 0.5 as “weakly correlated.”

#### Behavioral state determination

Trials were designated as stationary or running based on the mean forward wheel speed during the stimulus period of each trial, with a threshold of 2 cm/s as the threshold for running.

Pupil size and position were extracted from each frame using the native MATLAB function *imfindcircles*, and quantified by averaging all frames during the stimulus period on each trial. To designate large and small pupil trials, we first combined all stationary trials across both imaging sessions, found the median size of this pooled data, and labelled trials with a pupil size less than the median as “small pupil” and those with a pupil size greater than the median as “large pupil.”

#### Capture quantification

To assess capture on HTP+ cells, we analyzed widefield images of Alexa647 fluorescence collected immediately before the two-photon imaging experiment. In ImageJ, we created a circular ROI around the region of dTomato expression (**Figure S2E-F**), and measured mean fluorescence intensity within this ROI as well as 20-pixel perimeter around the ROI, to assess background fluorescence. We defined the Capture Index as:

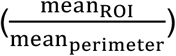

where values greater than 1 indicate enrichment of the DART ligands at the site of viral expression.

### Computational modeling

#### Model equations

We started from a four-population rate-based model, including pyramidal (E), PV (P), SST (S) and VIP (V) neuron populations ^10,11,25,26,34^. The firing rates of these populations (*r*_*E*_, *r*_*P*_, *r*_*S*_ and *r*_*V*_) obey standard rate equations

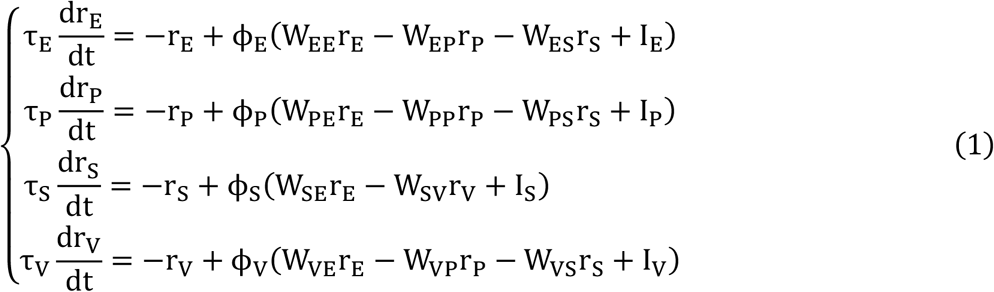

where W_AV_ is the strength of connections from population B to A, and I_A_, τ_A_, and ϕ_A_ are external inputs, time constant and transfer function (F-I curve) of population A. We used rectified-quadratic transfer functions for populations E and S (Rubin et al 2015), while for simplicity we used threshold-linear transfer functions for P and V populations:

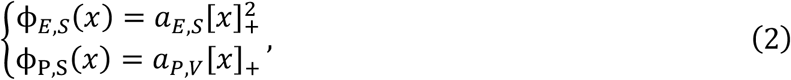

where [*x*]_+_ = 0 for x<0, [*x*]_+_ = *x* for x>0, while *a*_*E*,*S*_ and *a*_*P*,*V*_ are the gains for quadratic and linear transfer functions.

The influence of YM90K^DART^ is modeled as a decrease in the connection weight from Pyr neurons to SST cells as

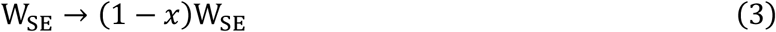

#### Reduction to a two population (E,S) model

To focus on the interactions between E and S cells, we simplified the four-population model into a two-population circuit composed of pyramidal cells and SST cells (**Figure 1A and S1**). In a steady state, we can derive from ***Equations 1***, the firing rates of P and V cells as a function of E and S cells exclusively:

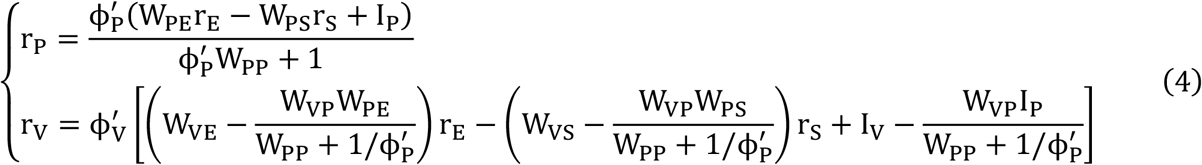

Combining ***Equations 1 and 4***, the firing rates of pyramidal cells obey

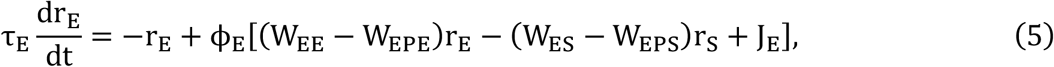

where 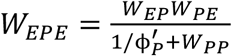 is the strength of the feedback of PV interneurons onto pyramidal cells, 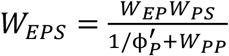 is the strength of the disinhibition of SST inhibition onto Pyr neurons through PV interneurons, and *J*_*E*_ is an effective external input to Pyr cells, defined as 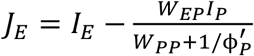 that includes feedforward inhibition from PV cells.

The firing rates of SST cells obey, respectively, in control group and DART group

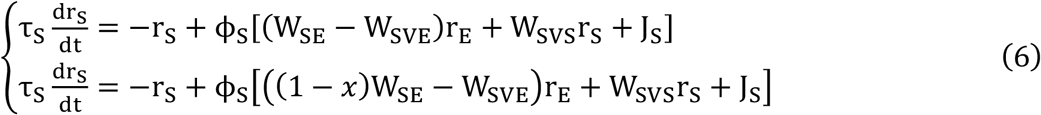

where 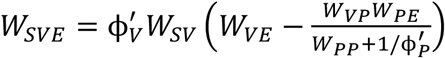 describes indirect effects of Pyr cells onto SST cells through VIP cells, 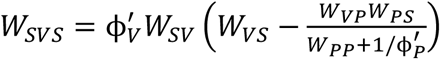 describes the strength of the feedback loop between VIP and SST cells, and *J*_*S*_ is an effective external input to SST cells, defined as 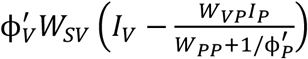 that includes overall inhibition from VIP cells.

#### Nullclines

The advantage of simplifying the model to two variables is that the dynamics of the model can be visualized on a 2-D plane spanned by the E and S rates. To get insight into the behavior of the model, it is useful to plot nullclines of the system, i.e. the curve on which the E rate is at equilibrium given r_*S*_ (the so-called r_*E*_ nullcline), and vice versa the curve on which the S rate is at equilibrium given r_*E*_ (the r_*S*_ nullcline). These nullclines are defined by setting the temporal derivatives of the rates to zero, i.e. 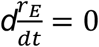 in ***Equation 5***, and 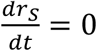 in ***Equation 6***. Fixed points of the network dynamics are then given by the intersections of these two nullclines. We first consider a simplified case where both E and S have linear transfer functions ϕ_*E*_ and ϕ_*S*_. In this case, the nullclines are given by:

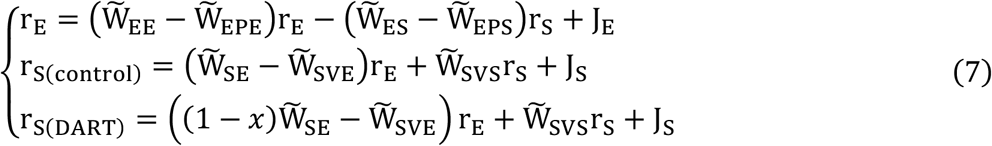

where 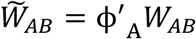 for all A,B=E,S, and 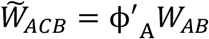 for all A,B=E,S and C=P,V. From ***Equation 7***, we find that the r_S_ nullcline increases monotonically with r_E_, with a slope that decreases in the DART condition, provided 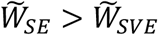 and 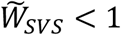

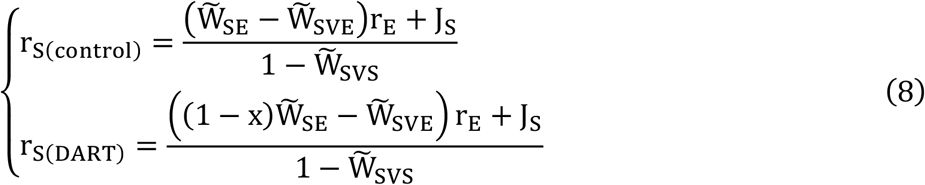

The r_E_ nullcline is given by:

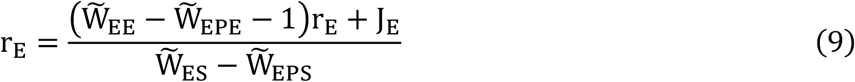

The sign of the slope of the r_E_ nullcline is determined by the sign of 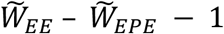 and 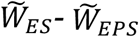. When 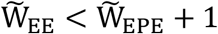 and 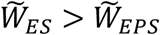, the slope of the r_E_ nullcline is negative (region *R*_*i*_). Thus, in this region, YM90K^DART^ leads to an increase in r_E_ and a decrease in r_s_ When 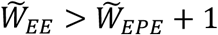 and 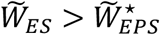, the slope of r_E_ nullcline becomes positive (region *R*_*ii*_). Thus, in this region, YM90K^DART^ leads to an increase of both r_E_ and r_s_. When 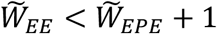 and 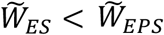, the slope of r_E_ nullcline is again positive (region *R*_*iii*_), but YM90K^DART^ leads to a decrease of both r_E_ and r_s_ (**Figure 1B-D**). The characteristics of each region can be summarized as follows:

| Region | slope of $r_E$ nullcline | Numerator | Denominator | YM90K<sup>DART</sup> effect |
| --- | --- | --- | --- | --- |
| $R_i$ | (-) | (-): $\tilde{W}_{EE} < \tilde{W}_{EPE} + 1$ | (+): $\tilde{W}_{ES} > \tilde{W}_{EPS}$ | ↓S ↑E |
| $R_{ii}$ | (+) | (+): $\tilde{W}_{EE} > \tilde{W}_{EPE} + 1$ | (+): $\tilde{W}_{ES} > \tilde{W}_{EPS}$ | ↑S ↑E |
| $R_{iii}$ | (+) | (-): $\tilde{W}_{EE} < \tilde{W}_{EPE} + 1$ | (-): $\tilde{W}_{ES} < \tilde{W}_{EPS}$ | ↓S ↓E |

#### Instability line

The stability of the fixed points of ***Equations 5***,***6*** can be determined by computing the eigenvalues of the Jacobian matrix of the system. In particular, a “rate” instability is reached whenever the Jacobian matrix has a zero eigenvalue, or equivalently *Det*(*J*) = 0 where *J* is the Jacobian matrix. This condition leads to

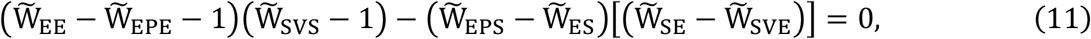

or equivalently

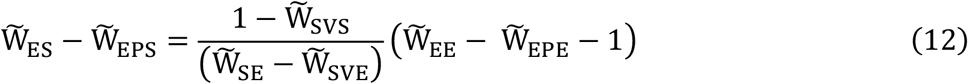

This line is plotted in **Figures 1** and **6. *Equations 5***,***6*** also potentially exhibit oscillatory instabilities in the ISN region, that depend on time constants in addition to effective weights. We checked that for parameters fitting the data, the model is stable with respect to such oscillatory instabilities. However, the model tends to develop damped oscillations in response to high contrast inputs, consistent with experimental observations in mouse visual cortex^56^.

#### Fitting procedure

The equations of the reduced two population model show that the fixed point of network equations depend only on five parameters involving the couplings: *W*_*EE*_ − *W*_*EPE*_, *W*_*ES*_ − *W*_*EPS*_, *W*_*SE*_, *W*_*SVP*_, *W*_*SVS*_. These equations also depend on *x*, the fractional reduction of AMPA receptor conductance by YM90K^DART^, and external inputs J_E_, J_s_. We used three variants of the model (Full, VIP, and Input; **Figure 6D** and **S6A-B**), that differ according to which parameters depend on state. In all models, external inputs depend on both contrast and state, and coupling strengths are independent of contrast. For both states and all contrasts, external inputs were constrained to produce the experimentally observed rates in control condition,

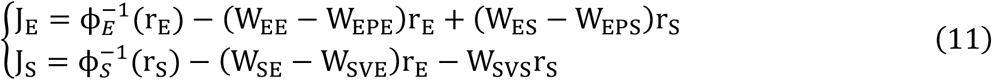

In the Full model, all coupling strengths depend on state. In the VIP model, all synaptic strengths are independent of state, but the gain of the VIP population 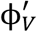 depends on state. We denote by *g* the ratio between VIP gain in running and stationary conditions. Note that this change only affects the effective weights that depends on VIP gain, i.e. *W*_*SVE*_ and *W*_*SVS*_. Finally, in the Input model, all weight parameters are fixed and independent of state. In all variants, *x* is a fixed parameter, independent of contrast and state. The value of *x* was set to 0.5, but we found that the minimum of the cost function C is independent of x, provided effective weights onto SST cells are varied accordingly (see below).

We defined a cost function C as

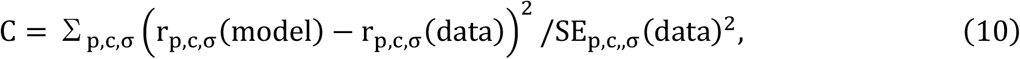

where the sum over *P* is a sum over populations (p = E, *S*), *c* = 25%, 50 %, 100 % is the contrast, and σ = stationary, running is the state. Note that in ***Equation 10*** only the YM90K^DART^ condition enters, since by construction all models in all conditions match the data perfectly in control conditions, provided the system converges to a fixed point. In some cases, the fixed point becomes unstable and the system converges to an oscillatory state, leading to a small discrepancy between model and data in control conditions. This happens in particular for the best fit ‘Input’ model at high contrast in running conditions (Figure S6B).

For each parameter set, modeled rates were obtained by simulating model equations. We then used the *differential_evolution* optimization algorithms from Python package *SciPy*.*optimize* to obtain the minimum of the cost function. We constrained the absolute value of all weight parameters to be smaller than 10, to avoid convergence to unrealistically large values of such parameters. For model selection, we used the Akaike Information Criterion (AIC)^58^. The optimal parameters found by this approach are shown in **Table 1** for the VIP model, and **Tables S1** and **S2** for the Full and Input models.

To show that the minimum of the cost function is independent of *x*, we first note that in control and YM90K^DART^ groups, Pyr influences SST through effective weights *A* (in control) and *B* (in YM90K^DART^),

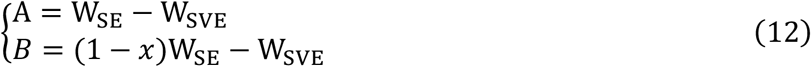

Once W_SE_ and W_SVE_ are found for a particular value of *x*, their values for arbitrary values of *x* can be obtained using

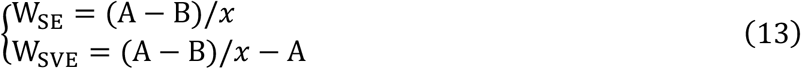

As *x* increases, W_SE_ and W_SVE_ decrease monotonically (**Figure S6C**). While W_*S*E_ is always positive (as it should be), W_SVE_ becomes negative for large enough *x*, which means that the indirect effect of Pyr→PV→VIP→SST disinhibitory pathway is stronger than the Pyr→VIP→SST inhibitory pathway.

## Supplementary Figure Legends

**Figure S1.**
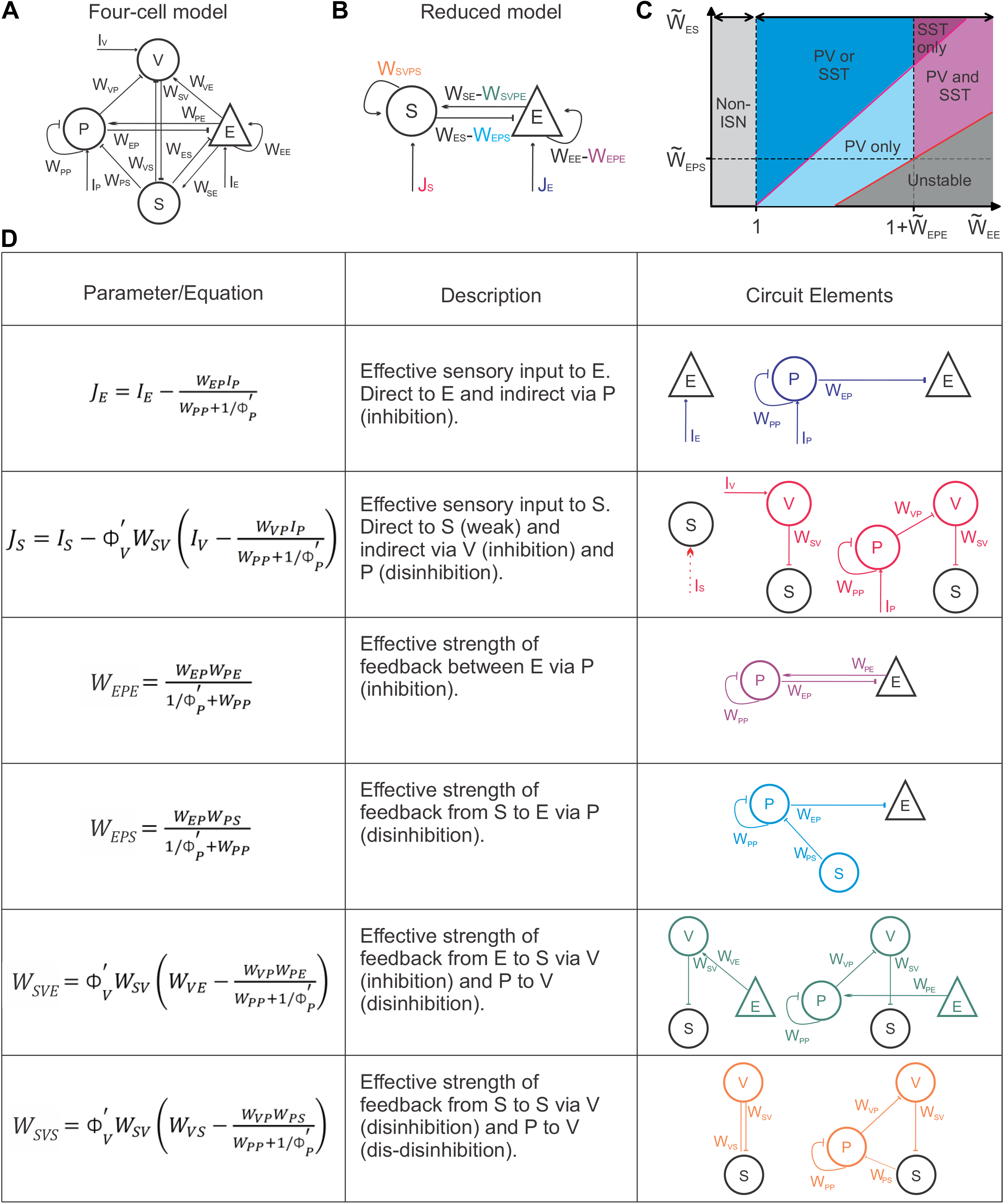
Definitions of connectivity weights in the reduced two-cell type model, related to Figure 1. (A) Schematic of the four-cell-type model with all input (I) and local (W) weights. (B) Schematic of reduced, two-cell-type model. *W*_*SE*_ and *W*_*ES*_ reflect direct connections between E and S cells; inputs (J) and other weights include connectivity of P and V cells. (C) Requirement for PV and SST cells in the space defined by 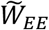 and 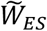. In the blue regions, PV cells are sufficient for stabilization; in the magenta regions, SST cells are required. (D) Table of equations, definitions and connectivity of inputs (J) effective weights (*W*_*EPE*_, *W*_*EPS*_, *W*_*SVE*_, *W*_*SVS*_). Colors in the connectivity diagrams correspond to weights in (B).

**Figure S2.**
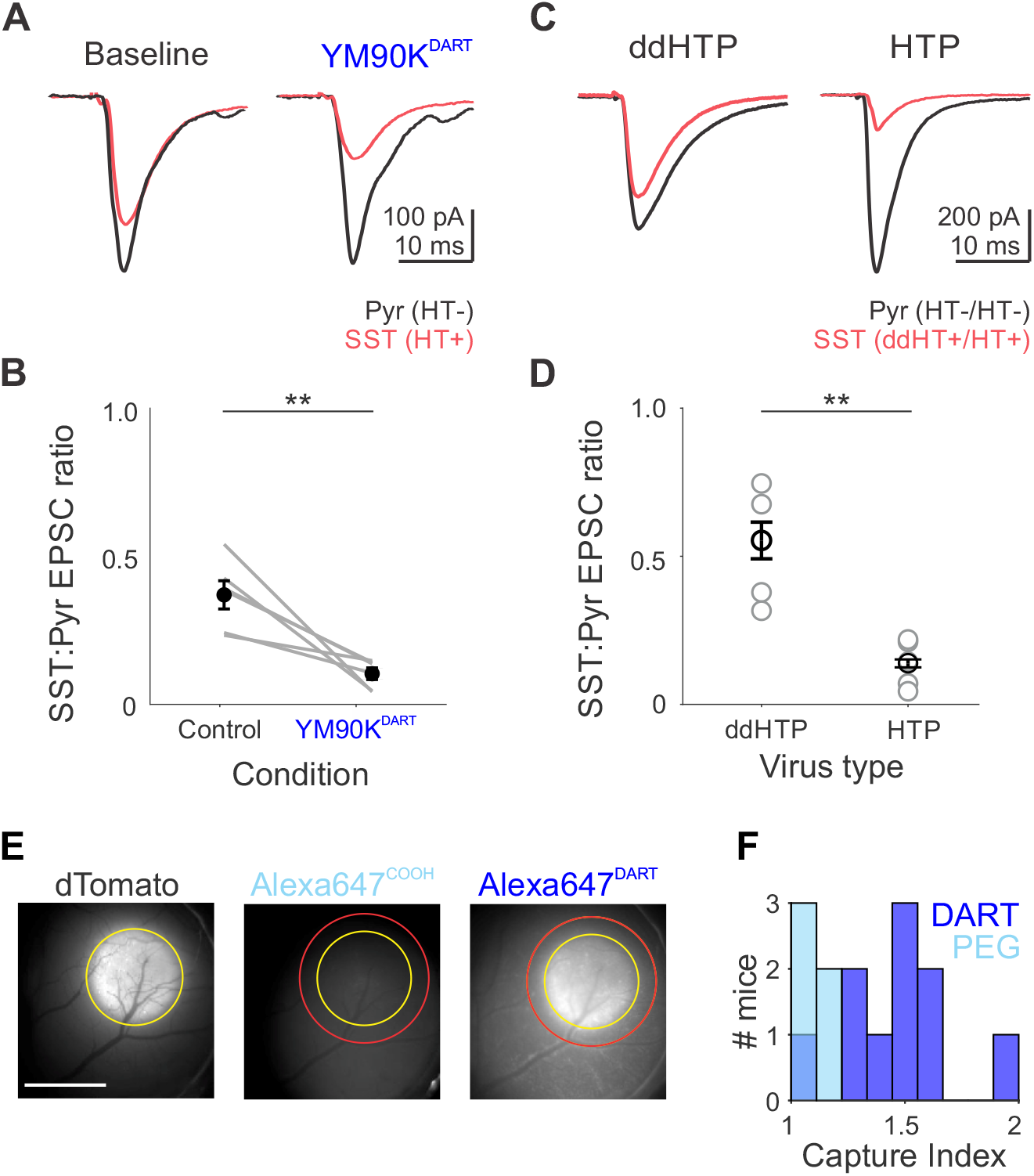
Selectivity of YM90K^DART^ antagonism and capture, related to Figure 2. (A) EPSCs in an example simultaneously recorded pair of SST (red) pyramidal (black) cells before (left), and after (right) application of YM90K^DART^ (300 nM) and Alexa647^DART^ (100 nM). (B) Summary of the ratio of SST to pyramidal EPSC amplitudes in control and YM90K^DART^. Grey lines connect pairs of cells (n = 6) recorded across conditions, black circles are the mean. Error bar is SEM across cell pairs. Paired t-test, p = 0.008. (C) EPSCs recorded in two example simultaneously recorded pairs of SST (red) pyramidal (black) cells following systemic infusion YM90K^DART^ (3 mM) and Alexa647^DART^ (0.3 mM) to the cerebrospinal fluid via the cisterna magna. The SST cell expresses either the non-binding ddHTP (left), or functional HTP (right). (D) Summary of the ratio of SST to pyramidal EPSC amplitudes for SST cells expressing either ddHTP (n = 4) or HTP (n = 5). Unpaired t-test, p = 0.003. (E) Example widefield images used to calculate the Capture Index. Left, dTomato expression was used to create an ROI (region of interest; yellow circle) around the HTP region. The ROI was applied to quantify intensity of either the non-binding Alexa647^COOH^ (middle) or Alexa647^DART^ (right) which were co-infused with YM90K^PEG^ or YM90K^DART^, respectively. A 20-pixel perimeter (red circle) was applied to measure background fluorescence. Scalebar = 200µM. (F) Distribution of Capture Index 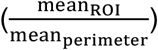 values for all YM90K^DART^ (dark blue, n = 10 mice) and YM90K^PEG^ experiments (light blue, n = 6 mice). * p < 0.05; ** p < 0.01.

**Figure S3.**
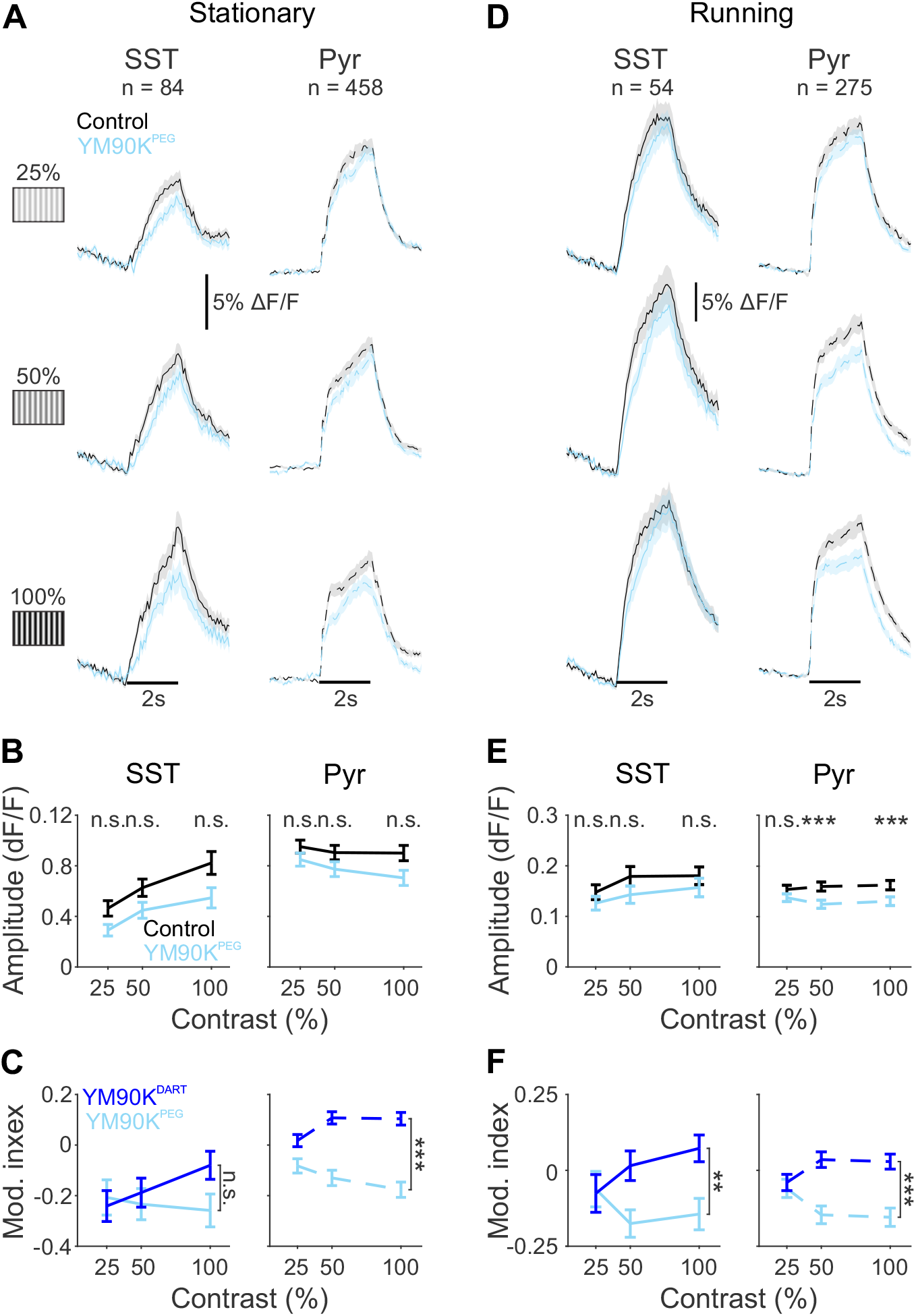
The effects of non-binding AMPAR antagonist YM90K^PEG^ and repeated imaging do not depend on contrast or behavioral state, related to Figure 3. (A) Grand average time courses for SST cells (left) and putative pyramidal cells (right) before (black) and after (light blue) YM90K^PEG^ during stationary epochs, at each contrast. Shaded error represents SEM across cells.(B) Mean response during stimulus period, for SST cells (left) and putative pyramidal cells (right) during stationary epochs, at each contrast. Error is SEM across cells. Two way ANOVA reveals a main effect for PEG within both SST (p = 0.001) and pyramidal (p = 0.003) cells; displayed significance refers to pair-wise Bonferroni-corrected t-tests between control and YM90K^PEG^ at each contrast. (C) Modulation index 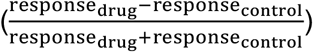 in SST cells (left) and putative pyramidal cells (right) following either YM90K^DART^ (blue) or YM90K^PEG^ (light blue). Significance refers to drug x contrast interaction from a two-way ANOVA, showing a trend toward facilitation by YM90K^DART^ for SST cells (p = 0.102), and a strong relative facilitation by YM90K^DART^ in pyramidal cells (p < 0.001). (D-F) Same as A-C, during running epochs, for the subset of cells with preferred-direction trials during running at all contrasts. For E, two way ANOVA reveals a main effect for PEG within both SST (p < 0.001) and pyramidal (p < 0.001) cells. For F, drug x contrast interaction shows robust relative facilitation by YM90K^DART^ for both SST cells (p = 0.005) and pyramidal cells (p < 0.001). Error is SEM across cells. n.s.-not significant; * p < 0.05; ** p < 0.01, *** p < 0.001

**Figure S4.**
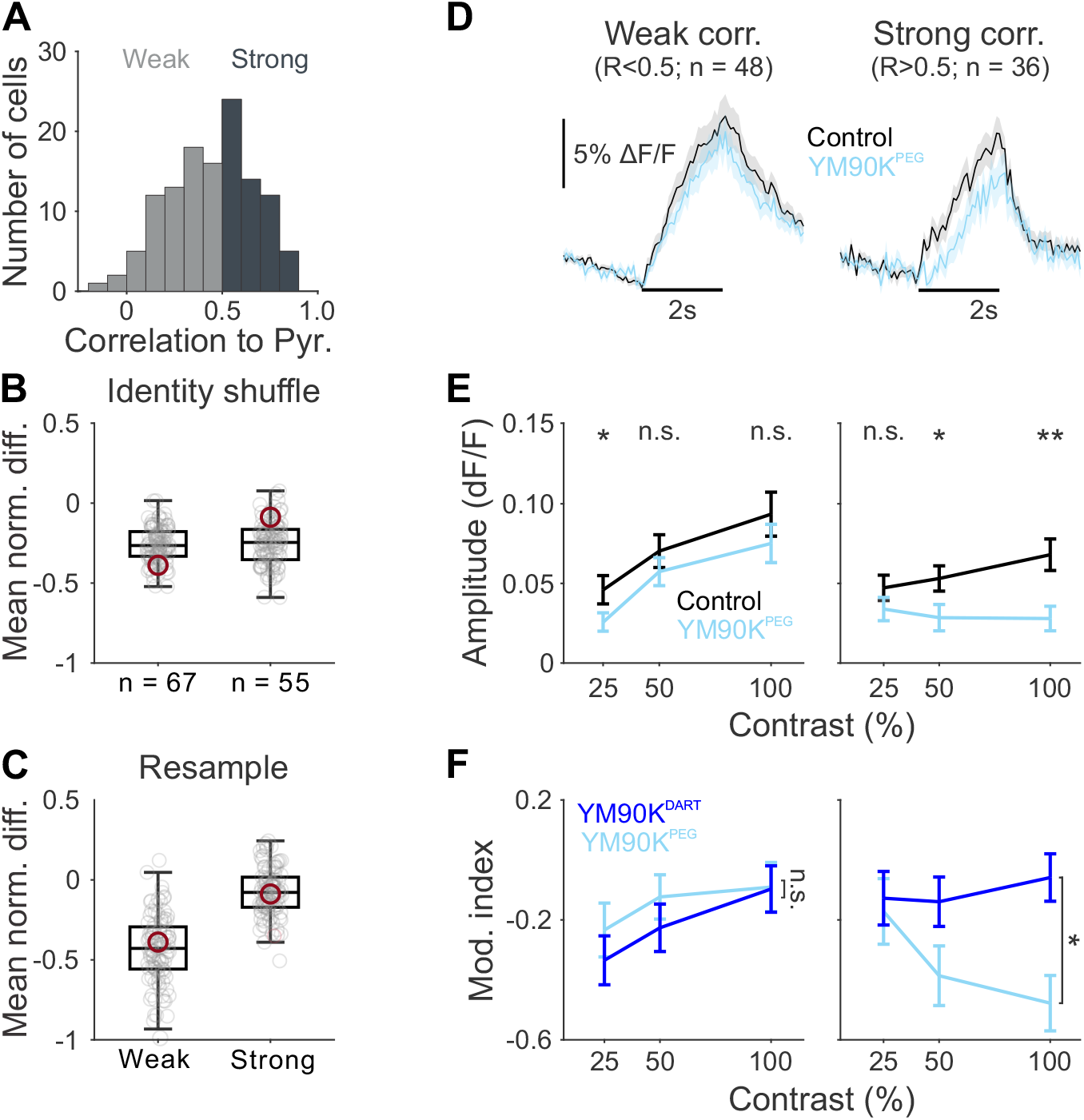
Correlation with the local network robustly and specifically predicts the effect of blocking AMPARs on SST cells, related to Figure 4. (A) Distribution of correlation coefficients for SST cells divided into weak (R < 0.5; light gray) and strong (R > 0.5; dark gray). (B) Distribution of mean normalized difference values of SST cells, when SST cell identity was shuffled across category (i.e. randomly sorted into mock “weak” and “strong” categories) 100 times. Each gray circle is the mean of one shuffle; box plots illustrate median, 25% and 75% quartiles. Maroon circles are the mean difference values with the correct identity assignment. Note that randomly separating cells into groups of these sizes does not produce differences between the groups on average. Cohen’s D for difference between groups = 0.083. (C) Same as (B), when SST cell identity was resampled with replacement within category 100 times. Cohen’s D for difference between groups = 1.743. (D) Grand average time courses for SST cells before (black) and after (light blue) YM90K^PEG^ separated into those weakly (R < 0.5) and strongly (R > 0.5) correlated to pyramidal activity, during stationary epochs in response to preferred-direction gratings at 50% contrast. Shaded error is SEM across cells. (E) Mean response during stimulus period, for SST cells weakly (left) or strongly (right) correlated to pyramidal activity, at each contrast. Two-way ANOVA reveals a main effect by YM90K^PEG^ in both the weakly correlated (p = 0.048) and strongly correlated (p = 0.004) SST cells; displayed significance refers to pair-wise Bonferroni-corrected t-tests between control and YM90K^PEG^ at each contrast. (F) Modulation index for weakly correlated (left) and and strongly correlated (right) SST cells following either YM90K^DART^ (blue) or YM90K^PEG^ (light blue), during stationary epochs at each contrast. Two-way ANOVA reveals no significant interaction of drug and contrast for weakly correlated cells (p = 0.862), but a significant interaction for strongly correlated cells (p < 0.022).

**Figure S5.**
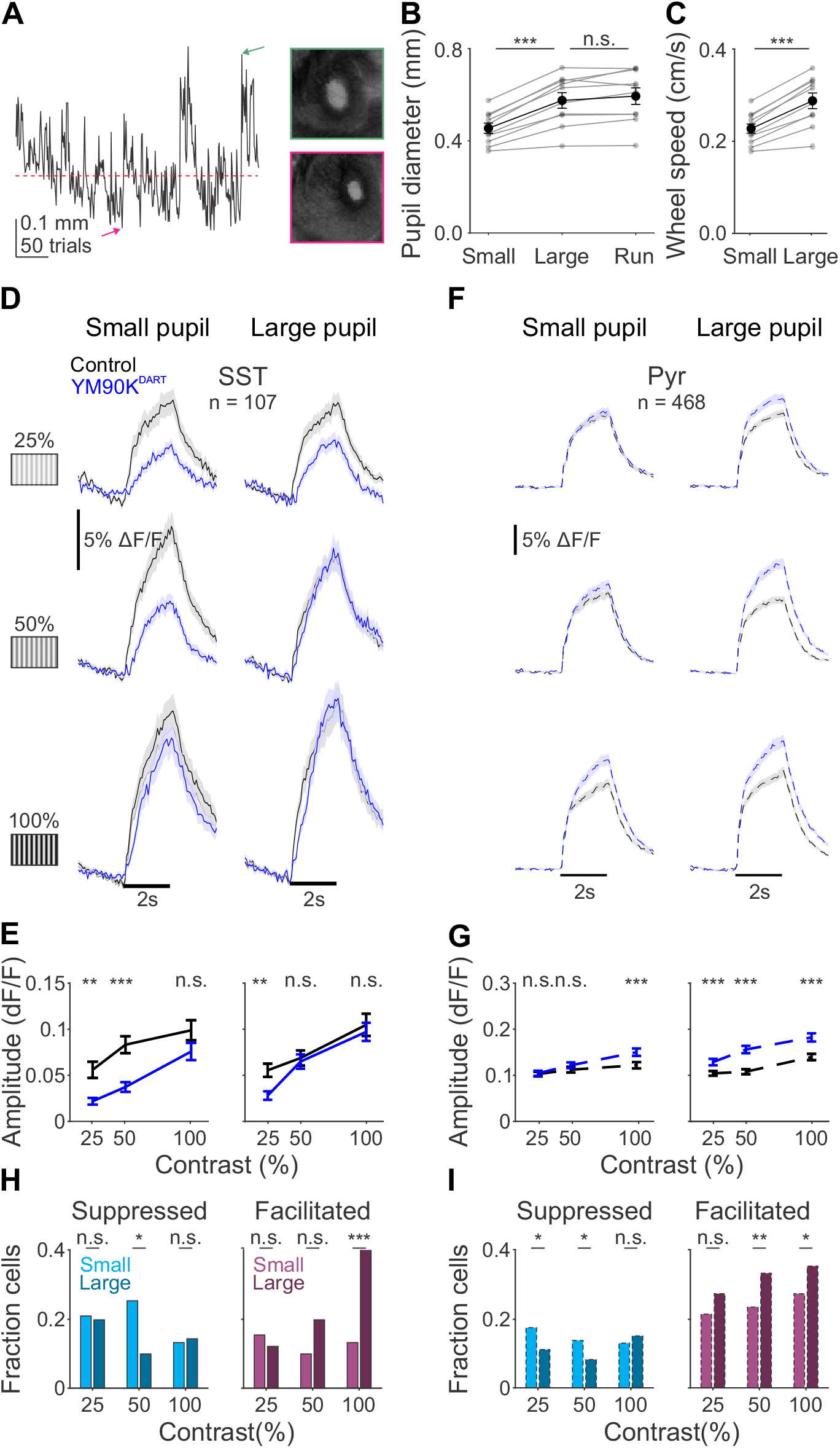
Arousal has similar effects to locomotion on the effect of blocking AMPARs on SST cells, related to Figure 5. (A) Left: timecourse of pupil sizes during stationary trials for an example experiment. Red line indicates median pupil size, used as threshold. Right: images of the pupil from representative large (top, green) and small (bottom, magenta) trials, from the times highlighted by colored arrows on the left. (B) Pupil diameter on small and large pupil trials during stationary epochs, and on running trials. Gray lines represent individual mice, black line represents mean. Error is SEM across mice. (C) Wheel speed for small and large pupil stationary trials. Note that the wheel speed threshold for locomotion is 2 cm/s. Gray lines represent individual mice, black line represents mean. Error is SEM across mice. (D) Grand average time courses for SST cells for small (left) or large (right) pupil trials, at each contrast before (black) and after (blue) YM90K^DART^ infusion. All cells are matched across pupil states and contrasts. Shaded error is SEM across cells. (E) Mean response during stimulus period for SST cells during small (left) or large (right) pupil trials, at each contrast. Error is SEM across cells. (F-G) Same as (A-B), for pyramidal cells. (H) Fraction of SST cells suppressed (left, cyan) or facilitated (right, magenta) by more than 1 std of their control response during small pupil (light) or large pupil (dark) epochs. (I) Same as H, for pyramidal cells.

**Figure S6.**
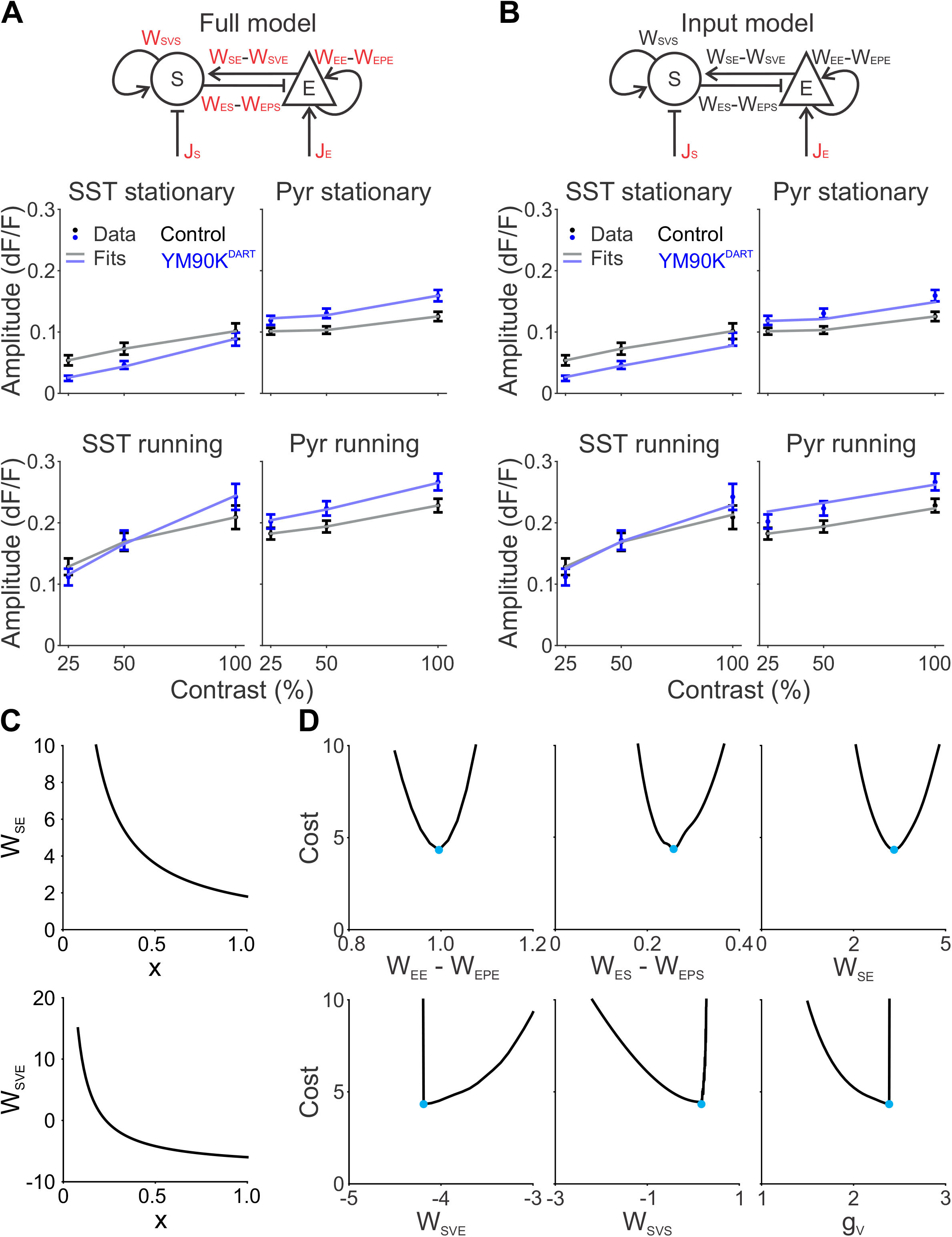
VIP model fits are superior to other models and robust to small changes in individual parameters, related to Figure 6. (A) Top, schematic of the Full model. Parameters in red are allowed to change across state. Bottom, empirical (dark data points, mean +/-SEM) and simulated (light lines) responses of SST (left) and pyramidal (right) cells to increasing contrast, in stationary (top) or locomotion (bottom) states in control (gray) and after YM90K^DART^ (blue). (B) Same as A, for the Input model. (C) Top, fit of *W*_*SE*_ as a function of *x*. Bottom, fit of for *W*_*SVE*_ as a function of x. (D) Fit cost for varying values of *W*_*EE*_ − *W*_*EPE*_, *W*_*ES*_ − *W*_*EPS*_, *W*_*SE*_, *W*_*SVE*_, *W*_*SVS*_, and *g* when *x* and other parameters are held constant. Cyan points are the fitted values. C and D are for the VIP model.

**Table S1.** Best fit parameters for “Full” V1 network model, related to Table 1. Effective connectivity weights are allowed to change across behavioral state but are held constant across contrast within state, while external inputs vary with stimulus contrast and state. Weights reflect the minimum cost found independently in the stationary and running states.

| *Parameters* | *Stationary* |  |  | *Running* |  |  |
| --- | --- | --- | --- | --- | --- | --- |
|  | 25% | 50% | 100% | 25% | 50% | 100% |
| $W_{EE} - W_{EPE}$ | 1.113 | | | 0.957 | | |
| $W_{ES} - W_{EPS}$ | 0.155 | | | 0.076 | | |
| $W_{SE}$ | 2.227 | | | 0.335 | | |
| $W_{SVE}$ | -2.933 | | | -0.959 | | |
| $W_{SVS}$ | 1.125 | | | 0.725 | | |
| $J_E$ | 0.176 | 0.179 | 0.188 | 0.211 | 0.215 | 0.218 |
| $J_S$ | -0.409 | -0.409 | -0.518 | -0.032 | -0.032 | -0.067 |

**Table S2.** Best fit parameters for “Input” V1 network model, related to Table 1. Effective connectivity weights are held constant across behavioral states, while external inputs vary with stimulus contrast and state.

| *Parameters* | *Stationary* |  |  | *Running* |  |  |
| --- | --- | --- | --- | --- | --- | --- |
|  | 25% | 50% | 100% | 25% | 50% | 100% |
| $W_{EE} - W_{EPE}$ | 0.959 | | | | | |
| $W_{ES} - W_{EPS}$ | 0.229 | | | | | |
| $W_{SE}$ | 4.948 | | | | | |
| $W_{SVE}$ | -10.000 | | | | | |
| $W_{SVS}$ | 0.582 | | | | | |
| $J_E$ | 0.195 | 0.201 | 0.215 | 0.230 | 0.240 | 0.250 |
| $J_S$ | -1.353 | -1.363 | -1.672 | -2.507 | -2.659 | -3.157 |

